# Master transcription factor binding sites constitute the core of early replication control elements

**DOI:** 10.1101/2023.10.22.563497

**Authors:** Jesse L. Turner, Laura Hinojosa-Gonzalez, Takayo Sasaki, Satoshi Uchino, Athanasios Vouzas, Mariella S. Soto, Abhijit Chakraborty, Karen E. Alexander, Cheryl A. Fitch, Amber N. Brown, Ferhat Ay, David M. Gilbert

## Abstract

Eukaryotic genomes replicate in a defined temporal order called the replication timing (RT) program. RT is developmentally regulated with potential to drive cell fate transitions, but mechanisms controlling RT remain elusive. We previously identified “Early Replication Control Elements” (ERCEs) necessary for early RT, domain-wide transcription, 3D chromatin architecture and compartmentalization in mouse embryonic stem cells (mESCs) but, deletions identifying ERCEs were large and encompassed many putative regulatory elements. Here, we show that ERCEs are compound elements whose RT activity can largely be accounted for by multiple sites of diverse master transcription factor binding (subERCEs), distinguished from other such sites by their long-range interactions. While deletion of subERCEs had large effects on both transcription and RT, deleting transcription start sites eliminated nearly all transcription with moderate effects on RT. Our results suggest a model in which subERCEs respond to diverse master transcription factors by functioning both as transcription enhancers and as elements that organize chromatin domains structurally and support early RT, potentially providing a feed-forward loop to drive robust epigenomic change during cell fate transitions.

## INTRODUCTION

DNA replication is an essential process that facilitates the faithful duplication of genetic information. In eukaryotes, chromosomes are replicated in a defined temporal order called the replication timing (RT) program. In mammals, changes in RT occur during development in units of 400-800 kilobases termed replication domains (RD), and changes in RT during development correlate with changes in transcriptional programs (Rivera-Mulia *et al*, 2019a; Vouzas & Gilbert, 2023). Defects in RT are correlated with dysregulation of genes and genome instability, observed in several diseases and cancers, and associated with defects in chromosome condensation and sister chromatid cohesion (Blumenfeld *et al*, 2017; Heskett *et al*, 2020; Platt *et al*, 2018; Rivera-Mulia *et al*, 2017; Rivera-Mulia *et al*, 2019b; Sasaki *et al*, 2017). Recently, it was revealed that RT perturbations result in immediate epigenetic disturbances (Klein *et al*, 2021), demonstrating that RT is essential to maintain the epigenome.

We previously demonstrated (Sima *et al*, 2019) that early RT of several RDs in mouse embryonic stem cells (mESCs) is controlled by discrete, cooperative, and redundant chromosome segments termed early replication control elements (ERCEs). ERCEs are required for early RT, transcription, euchromatic compartmentalization, and Topologically Associating Domain (TAD) organization of their entire domain. However, the deletions identifying ERCEs spanned tens of kilobases and multiple discrete regions with active chromatin signatures, leaving open the possibility that they may contain several different types of control elements. ERCEs contain enhancer histone marks, the histone acetyltransferase CBP/P300, multiple Oct4/Sox2/Nanog co-bound (OSN) sites, transcription start sites (TSS), and they interact in a CTCF-independent manner. Changes in cooperative binding of master transcriptional regulatory network (TRN) transcription factors (TFs) have been shown to strongly correlate with changes in RT during differentiation (Rivera-Mulia *et al*., 2019). In budding yeast, TFs Fkh1/Fkh2 mediate early RT through spatial clustering of early origins, independent of their transcriptional activity (Fang *et al*, 2017; Knott *et al*, 2012; Ostrow *et al*, 2017). Together, these results suggest the hypothesis that ERCE activity may reside within their resident master TF co-binding sites.

To test this hypothesis, we performed a series of finer deletions to delineate the locations of ERCE activity. We found that OSN sites contributed to RT activity in all 3 regions, in some contexts completely accounting for ERCE activity. By contrast, deletion of all 3 TSSs that resided within the prior deletions eliminated nearly all transcription in the domain without loss of early replication. Thus, early replication can be maintained in the face of of severe reductions in transcriptional activity. We find that OSN sites are also bound by many diverse master TFs and are unlikely to derive their RT activity from the OSN TFs themselves designate them as “subERCEs”, novel functional elements that can both enhance transcription but also play non-transcriptional roles in RT. subERCEs do not overlap efficient replication origins, and a large deletion encompassing most of the replication initiation activity in the domain did not affect RT, demonstrating that subERCEs regulate RT domain-wide, independent of where replication initiates. Together with our prior demonstration of their 3D interactions, supported by more recent HiChIP and Micro-C data, we propose that ERCEs are compound elements that create a sub-nuclear environment that is favorable for early firing of potential replication origins located throughout the domain. Notably, our dissections highlight the robustness and cooperativity of ERCEs, which function as redundant compound cis-regulatory elements to maintain early replication.

## RESULTS

### Functional dissection of the Dppa2/4 replication domain using allele-specific deletions

The ∼400kb murine “Dppa domain” contains three active genes in mESCs, *Dppa2*, *Dppa4*, and *Morc1* (**Supplemental Fig. 1**). *Dppa2* and *Dppa4* are markers of pluripotency (Maldonado-Saldivia *et al*, 2007) and *Morc1* is essential for spermatogenesis (Inoue *et al*, 1999), although it is also highly expressed in ESCs (Sima *et al*., 2019). The Dppa domain corresponds to a TAD whose boundaries align with Hi-C A/B compartment boundaries and are demarcated by convergent CTCF binding sites that also are bound by cohesin (Sima *et al*., 2019). Our previous study identified three segments with ERCE activity in the domain, designated A, B and C (**Supplemental Fig. 1A-B**). These ERCE-containing segments interact and contribute to TAD strength in a CTCF-independent manner (Sima *et al*., 2019). ERCEs, but not CTCF protein or the CTCF-bound TAD boundaries, were necessary to maintain interactions with the A compartment and early replication. The Dppa domain resides in the nuclear interior but is flanked on both sides by late-replicating lamina-associated domains (Takebayashi *et al*, 2012a) and deletion of the ERCE-containing segments causes a movement of the Dppa domain toward the nuclear lamina (Brueckner *et al*, 2020).

During differentiation, the Dppa domain undergoes a domain-wide shift from early to late RT (**Supplemental Fig. 1Bi**), concomitant with a switch from the A compartment to the B compartment and silencing of genes throughout the domain (Hiratani *et al*, 2010; Hiratani *et al*, 2008; Sima *et al*., 2019; Takebayashi *et al*, 2012b). These changes occur in all three germ layers, thus the Dppa domain is a pluripotency-specific early replicating domain (Hiratani *et al*., 2010; Hiratani *et al*., 2008).

Our studies of *cis*-acting elements in the Dppa domain have been greatly facilitated by the use of mESCs derived from F1 *castaneusXmusculus* hybrid mice (F121-9). These two subspecies are separated by 500,000 years and harbor single nucleotide polymorphisms (SNPs) at, on average, every 150bp, facilitating distinction between paternal and maternal homologues (Rivera-Mulia *et al*, 2018). This has allowed us to make heterozygous manipulations in one allele (red lines in all figures) while retaining the WT allele in the same datasets (black lines in all figures) (**Supplemental Fig. 1C**). **Supplemental Figure 1Bii-viii** show the results of single, double, and triple ERCE deletions that were previously published (Sima *et al*., 2019).

**FIGURE 1.**
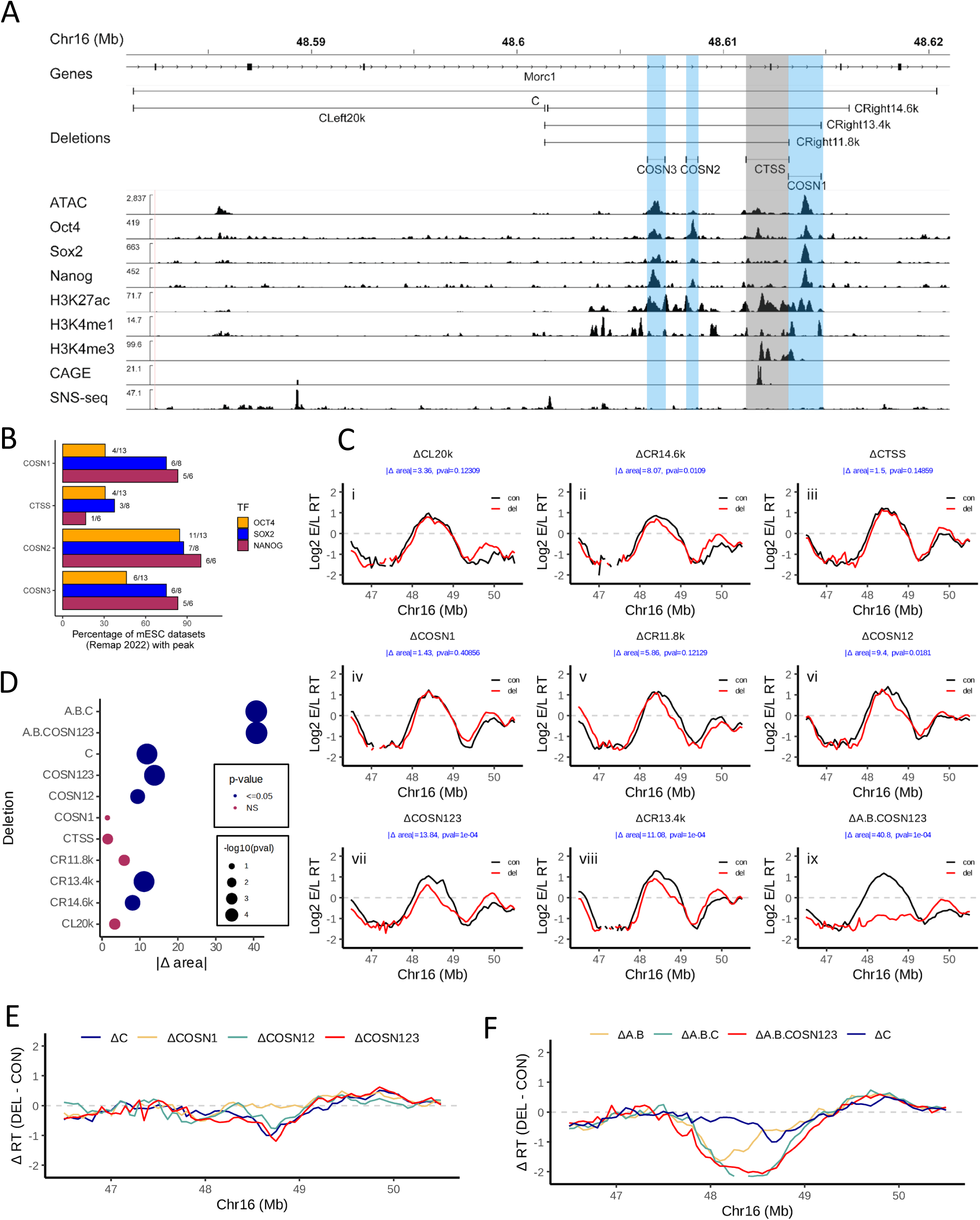
Genetic dissection of the C element implicates OSN binding regions as central to ERCE RT activity. A) IGV browser tracks of the Dppa domain, zooming into the C element. Positions of OSN and TSS are indicated in blue and grey, respectively. All deletions of the C element are represented by horizontal bars above the data tracks. B) Percentage of Remap2022 mESC Oct4, Sox2, and Nanog ChIPseq datasets with peaks that overlap the deletions generated in the C element. C) RT profiles in the Dppa domain of cell lines harboring allele-specific deletions (red) compared to the homologous unmodified allele (black) in C that are shown in panel A, averaged across replicates (independent CRISPR clones and technical replicates). Individual replicate experiments are shown in Supplemental Fig. 3. Δarea and pval, calculated as in Supplemental Figure 2, are shown in blue font in each RT panel. D) |Δ area| between the RT profile curves for the deletions shown in C and the significance of their delay in RT compared to the corresponding WT alleles (see Materials and Methods) . E-F) Comparison of RT changes [ΔRT (DEL-CON)] in the Dppa domain (E) comparing ΔC to sequential deletions of COSN elements (Sima et al., 2019) and COSN123, and (F) comparing ΔA.B.C (Sima et al., 2019) to ΔA.B.COSN123 with ΔAB and ΔC as comparators.

Given that we expected a series of smaller deletions to have more subtle effects on RT than our original deletion series, we took advantage of our hybrid system to develop a rigorous statistical method to determine the RT change for each deletion with respect to its corresponding internal control (WT) allele. This approach compares the area between two RT curves of the Dppa domain (WT vs deletion; **Supplemental Fig. 1C**) to area from similarly-sized random regions across the genome to test whether the observed delay for Dppa domain is significantly larger than expected by chance (**Supplemental Fig. 1D, Supplemental Figure 2A-B**; see **Materials and Methods**). Upon test-ing this approach on our previously published deletions, we observed statistically significant differences for ΔABC and all pairwise deletions as expected (**Supplemental Fig. 1D**). Among the single deletions, statistical significance was reached by B and C deletions but not for A deletion, which is consistent with visual patterns of RT curves (**Supplemental Fig. 1D**). These results demonstrate the effectiveness of our new quantitative approach, which becomes critical to assess the impact of smaller deletions while incorporating data from multiple deletion clones or replicates (**Supplemental Figure 2**; see **Materials and Methods**). These numbers are provided within each deletion panel in all figures throughout the manuscript (blue font; Δarea and pval in each RT panel). **Supplemental Fig. 3** shows all of the individual replicates from independent CRISPR-mediated deletion clones we generated for this study or re-analyzed from Sima et. al. **Supplemental Fig. 2B** compares the |Δarea| and pval for all of these same deletions. **Supplemental Tables 1, 2**, and **3** contain the sgRNA/primer sequences, coordinates of deletions as determined by PCR/Sanger sequencing/NGS analysis, and the genotypes of **65** cell lines, respectively.

### Cooperative Contributions of 3 OSN Sites Maintain Activity of ERCE C

We first performed a series of deletions in the 39kb ERCE C, the ERCE deletion with the largest effect on RT when deleted alone (**Supplemental Fig. 1B**). ERCE C contains 3 OSN sites (COSN3, COSN2, COSN1, 5’ to 3’) and one annotated intragenic TSS for Morc1 ∼135kb downstream from the start of the gene (CTSS) (**Fig. 1A-B**). To ensure that our classification of OSN sites was supported by multiple independent ChIP experiments, and to provide a quantitative assessment of the strength of each OSN site, we interrogated the Remap2022 database for mESC datasets (Hammal *et al*, 2022) and quantified the percentage of the available datasets for each TF that contained a peak overlapping our targeted deletions. This analysis revealed that OSN binding activity is detected at CTSS in some available datasets but much less reproducibly than the sites we refer to in this manuscript as OSN sites (**Fig. 1B**). Since the right half of C contains all active regulatory marks (**Fig. 1A and Supplemental Fig. 1A**) and an annotated mESC super-enhancer (Dowen *et al*, 2014; Khan & Zhang, 2016), we began by deleting the right and left halves of C. Deletion of the left half (ΔCL20k) had no effect while deletion of the right half (ΔCR14.6k) caused a significant delay that was reproducible in two independent deletion clones (**Fig. 1Ci and Fig. 1Cii; Supplemental Fig. 3 xxvi and xxxii; Fig. 1D).** We next focused our attention on the OSN and TSS sites in the right half of C. We deleted the TSS (ΔCTSS; **Fig 1Ciii**), the OSN site downstream of the TSS (ΔCOSN1; **Fig. 1Civ**), as well as a deletion of COSN2, CTSS and COSN3 that leaves COSN1 intact (ΔCR11.8k; **Fig. 1Cv; Supplemental Fig. 3xxx**), none of which resulted in a statistically significant change in RT compared to WT (**Fig. 1D**). These results suggested that ERCE C may consist of multiple cooperative elements, so we deleted COSN2 or both COSN2 and COSN3 in the ΔCOSN1 cell line, generating ΔCOSN12 and ΔCOSN123 (**Fig. 1C-vi and vii; Supplemental Fig. 3 xxviii and xxix**). ΔCOSN12 resulted in a significant RT delay (**Fig. 1D**), while deletion of all three OSNs (ΔCOSN123) gave an effect similar to deletion of the entire C element (**Fig. 1D-E; Supplemental Table 4**). We also generated a larger deletion that spans all 3 OSNs (ΔCR13.4k), which also resulted in a delay similar to ΔCOSN123, and ΔC (**Fig 1C-viii; Supplemental Fig. 3xxxi; Fig.1D; Supplemental Table 4**). In summary, all deletions that remove all three OSNs (ΔCR14.6k, ΔCR13.4k, ΔCOSN123) significantly delay RT (**Fig. 1D**), and these delays are not significantly different from that of ΔC (**Supplemental Table 4)** or from each other (**Supplemental Table 4)**. Interestingly, ΔCOSN12 also significantly delayed RT (**Fig. 1D**) while ΔCR11.8k (which deletes COSN23 and CTSS) did not significant delay RT (**Fig. 1C-D**), suggesting that not all combinations of OSN deletions are equivalent and that some OSNs play larger roles than others in different contexts.

Since we were only able to recover one clone of ΔCONS123 with all three deletions in the same allele, we performed 4 independent Repli-seq experiments starting from separate cultures of this cell line (**Supplemental Fig. 3xxix**). This provided more statistical power, and also allowed us to accumulate sufficient reads to verify that these were the only deletions in the Dppa locus throughout both alleles of this clone (**Supplemental Table 3**), detectable as missing sequences from one allele in the Repli-seq data. To evaluate whether ΔCONS123 resembles ΔC in other contexts, we introduced the full A and B deletions from our previous work into the ΔCOSN123 background, giving rise to cell line ΔA.B.COSN123 (**Fig. 1Cix; Supplemental Fig. 3iv**), which fully delayed RT reminiscent of and not significantly different from ΔA.B.C (**Fig. 1D-F**). We thus conclude that ERCE activity within the “C” element can be accounted for by the three OSN cobound sites.

### OSN sites contribute to the ERCE activity of the A and B elements

ERCE A, initially discovered with a 3.5kb deletion, contains one OSN site (AOSN) and one active TSS (ATSS) for the Dppa2 gene that also overlaps an OSN binding site (**Fig. 2A-B**). Because ΔA had no effect on RT on its own (**Supplemental Fig. 1B and D**), and because we had shown that ΔCOSN123 accounts for ΔC, we dissected the “A” element in the ΔCOSN123 background. Loss of the full activity of AOSN and ATSS is expected to delay RT similar to ΔA.C. First, we introduced either ΔAOSN or ΔA (removing both AOSN and ATSS) into the ΔCOSN123 cell line, creating ΔAOSN.COSN123 and ΔA.COSN123 (**Fig. 2Ci-ii and Supplemental Fig.3xvi and xi**). ΔA.COSN123 significantly delays RT relative to ΔCOSN123 but does not completely recapitulate ΔAC (**Fig. 2D and Supplemental Table 4**), suggesting that the OSN sites in C do not recapitulate the effect of the full C deletion when B is still intact. However, ΔA.COSN123 did cause a significantly greater delay compared to ΔAOSN.COSN123 (**Supplemental Table 4**). In fact, ΔAOSN.COSN123 had no additional contribution to the delay in RT relative to ΔCOSN123 alone (**Fig. 2E, Supplemental Fig. 3, and Supplemental Table 4**). Since ΔA is already relatively small, and since ΔATSS on its own does not delay RT in other contexts (discussed below), it is likely that AOSN and ATSS (which contains an OSN) are cooperative for maintaining RT at ERCE A.

**FIGURE 2.**
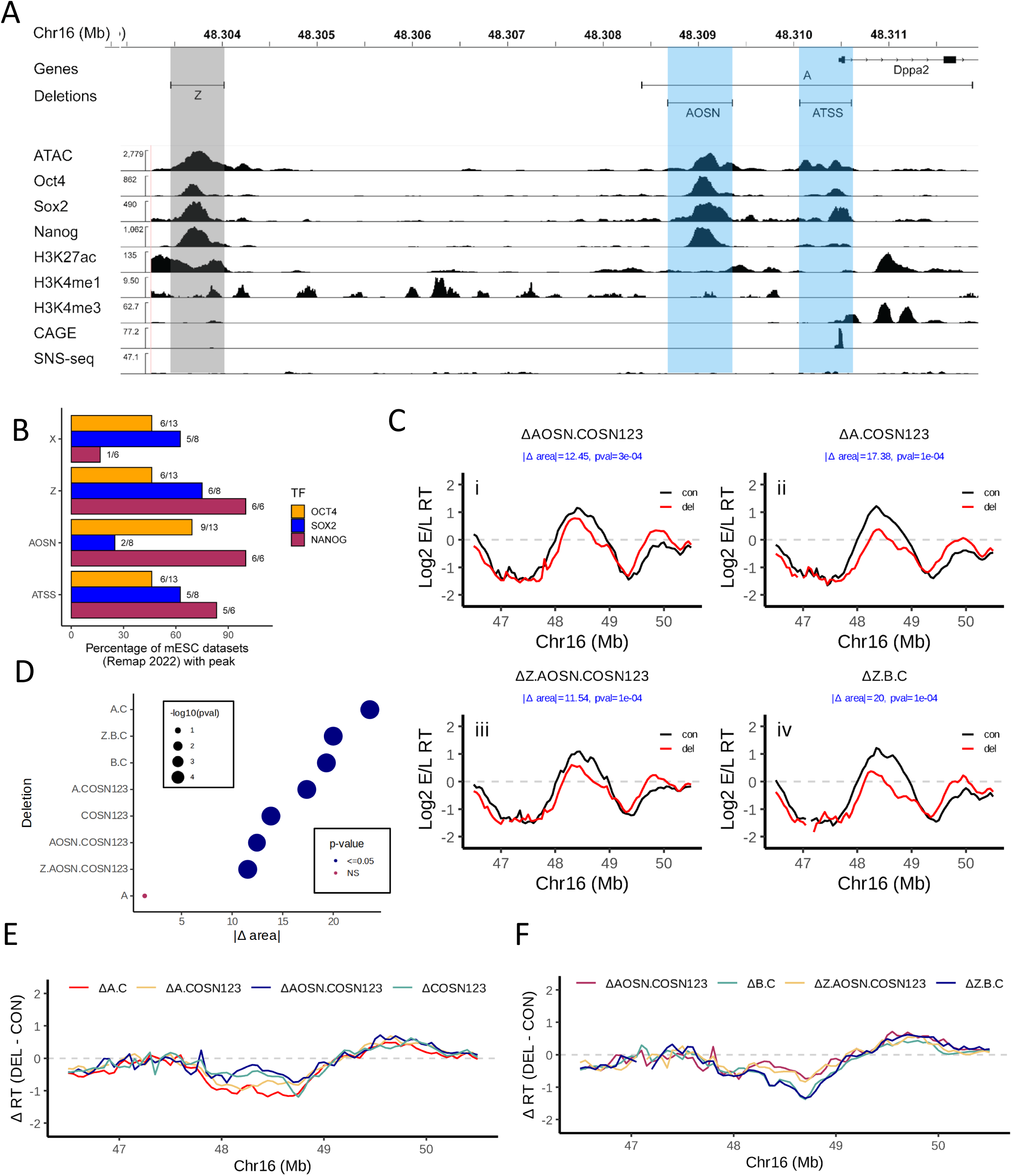
OSN sites contribute to the ERCE activity of the A element A) IGV browser tracks as in Figure 1A, zooming into the A element. Deletions The pvalue box is low resolution and illegible. of the A element are demarcated at the top by horizontal bars above the data tracks. Sites harboring activity are highlighted in blue while sites without activity (Z) are highlighted in grey. B) Percentage of Remap2022 mESC Oct4, Sox2, and Nanog ChIPseq datasets with peaks that overlap the deletions generated in the A element as well as an additional upstream site (X, shown in Supplemental Figure 1) that was shown to have no activity in our previous work (Sima et al., 2019). C) RT profiles of cell lines harboring deletions (red) compared to the homologous unmodified allele (black) in and near ERCE A, presented as in Figure 1C. D) |Δ area| between the RT profile curves of the deletions shown in C and the significance of their delay in RT compared to the corresponding WT alleles (see Materials and Methods). E-F) Comparison of RT changes [ΔRT (DEL-CON)] in the Dppa domain (E) comparing the deletion of A and AOSN in the ΔCOSN123 and ΔC backgrounds, and (F) comparing deletion of Z in the ΔAOSN.COSN123 and ΔB.C backgrounds.

Given our finding of the importance of OSN sites in the C element, we noted that upstream of the A element is another OSN (Z) site that is outside of the original A deletion. Deletion of Z in the context of either ΔAOSN1.COSN123 or ΔB.C, producing ΔZ.AOSN.COSN123 and ΔZ.B.C (**Fig. 2Ciii and iv and Supplemental Fig. 3xxxix-xl**), respectively, did not delay RT relative to their parental backgrounds (**Fig. 2D and F and Supplemental Table 4**). Consistent with its absence from ERCE A and its presence in ΔA.B.C, which is late replicating, we conclude that Z is neither necessary nor sufficient for ERCE activity, reminiscent of a strong OSN-containing region we previously designated as “Y” (**Supplemental Fig. 1,** (Sima *et al*., 2019). Since both Y and Z remain in-tact in the very late replicating ΔA.B.C and ΔA.B.COSN123 deletions, the presence of these two OSNs is not sufficient to advance RT. For this reason, we designate the compound elements within ERCEs that contribute to RT as “subERCEs”. It will be important to understand what essential component of subERCEs is missing from the Y and Z OSN binding sites (see Discussion).

ERCE B (∼45 kb in size) contains two OSN sites, one ∼8kb upstream (BOSN1) and another ∼4kb downstream (BOSN2) of the TSS for the large (200kb) Morc1 gene (BTSS) (**Fig. 3A-B**). BTSS does not contain any detectable OSN binding activity (**Fig. 3 A-B**). Since, like ERCE A, ΔB has a small effect on RT on its own, we dissected the “B” element in the context of other deletions. We first deleted BOSN1 in the context of ΔCOSN123 and ΔA.COSN123 to produce ΔBOSN1.COSN123 and ΔA.BOSN1.COSN123, which produced delays with no statistically significant difference compared to their parental backgrounds (**Fig. 3Ci and ii**, **Fig. 3D, Supplemental Fig. 3xxi and v, and Supplemental Table 4**). We then deleted BOSN2 in these contexts, producing ΔBOSN12.COSN123 and ΔA.BOSN12.COSN123 (**Fig. 3Ciii-iv and Supplemental Fig.3xxi and vii**). ΔBOSN12.COSN123 was statistically indistinguishable from ΔB.C and significantly more delayed than ΔBOSN1.COSN123, demonstrating that, in the absence of COSN123, deletion of BOSN12 can account for the B ERCE RT activity. However, while ΔA.BOSN12.COSN123 was significantly more delayed than ΔA.BOSN1.COSN123 (**Figure 3D and Supplemental Table 4**), ΔA.BOSN12.COSN123 was still significantly less delayed than either ΔA.B.C or ΔA.B.COSN123 (**Fig. 3D-E and Supplemental Table 4**), indicating that something else is contributing to the promotion of early replication in the absence of all OSN sites with ERCE activity. We also generated the BOSN deletions in the ΔAOSN.COSN123 background, producing ΔAOSN.BOSN1.COSN123 and ΔAOSN.BOSN12.COSN123 (**Figure 3Cv and 3Cvi and Supplemental Fig.3xiii-xiv**). Consistent with a role for BOSN2 in the activity of ERCE B, ΔAOSN.BOSN12.COSN123 but not ΔAOSN.BOSN1.COSN123, was significantly delayed compared to ΔAOSN.COSN123 (**Supplemental Table 4**). Surprisingly, however, ΔAOSN.BOSN12.COSN123 was not significantly more delayed than ΔAOSN.BOSN1.COSN123 (**Supplemental Table 4**) despite the large change in Δarea (12.7 vs. 19), but did show a change in shape of the RT curve that resembles ΔA.BOSN12.COSN123 (**Fig. 3Cii-vi**). Altogether, our results implicate the OSNs are important for B ERCE activity, but show that their ability to account for B ERCE activity is dependent on the presence of components of the A element via cooperativity relationships we do not yet understand.

**FIGURE 3.**
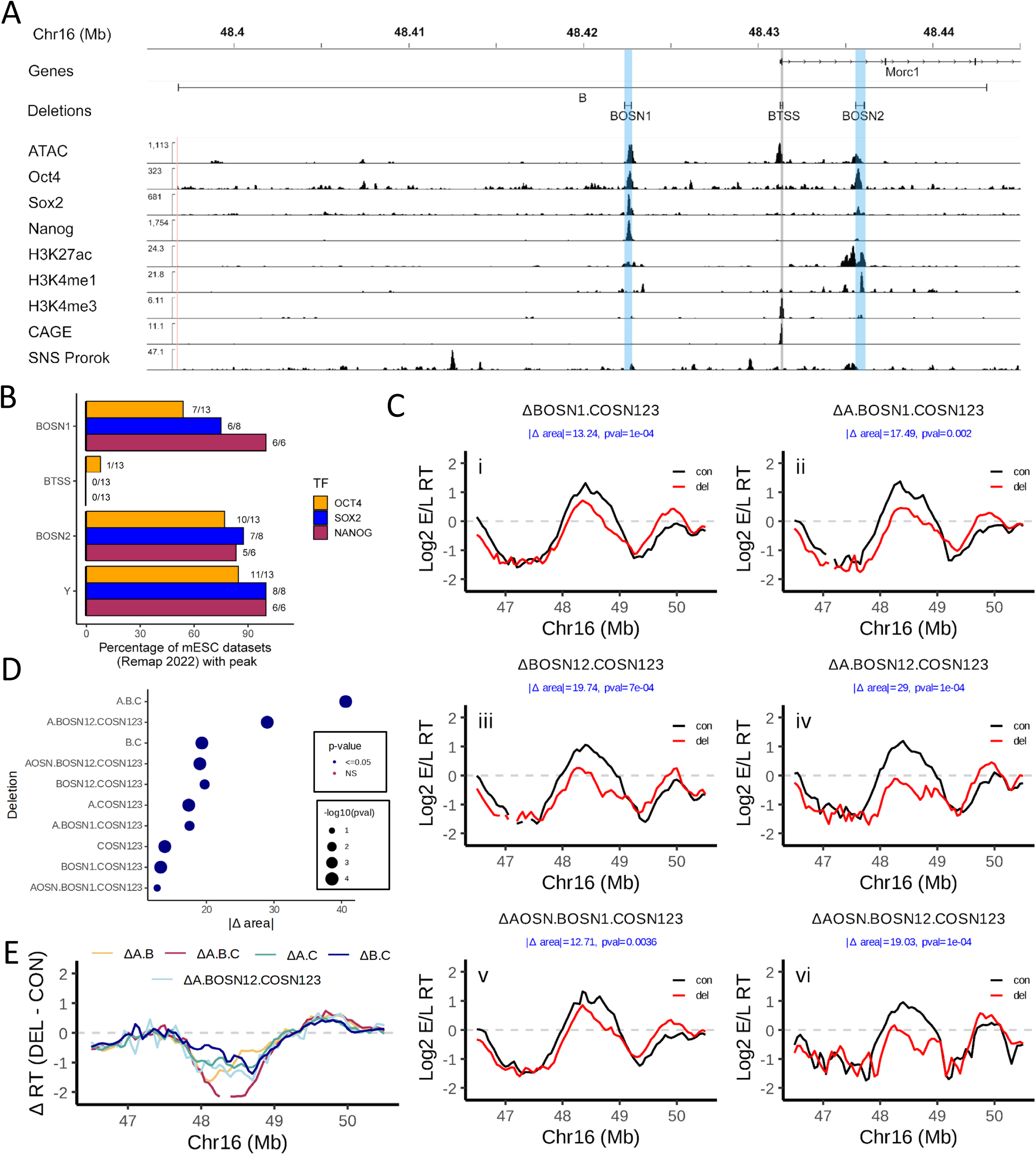
OSN sites contribute to the ERCE activity of the B element A) IGV browser tracks as in Figures 1A and 2A, zooming into the B element. Deletions of the B element are demarcated at the top by horizontal bars above the data tracks. OSNs are highlighted in blue while the TSS is highlighted in grey. B) Percentage of Remap2022 mESC Oct4, Sox2, and Nanog ChIPseq datasets with peaks that overlap the deletions generated in the B element as well as an additional site 12 kb downstream of B (Y, not shown in panel A) that was shown to have no activity in our previous work (Sima et al., 2019). C) RT profiles of cell lines harboring deletions (red) compared to the homologous unmodified WT allele (black) of sites within ERCE B presented as in Figure 1C. D) |Δ area| between the RT profile curves of the deletions shown in C and the significance of their delay in RT compared to the corresponding WT alleles (see Materials and Methods). E) Comparison of RT changes [ΔRT (DEL-CON)] in the Dppa domain comparing ΔA.BOSN12.COSN123 to ΔA.B.C and pairwise ERCE deletions.

### Early RT can be maintained in the absence of most transcription

In our previous study, we reported that a triple deletion of A, B and C led to silencing of transcription within the Dppa domain (Sima *et al*., 2019). However, it remained to be determined whether ERCE RT activity could be uncoupled from their transcription activity. This is particularly important in light of the complex effects of deletions that contained TSSs on early replication and transcription (Sima *et al*., 2019). To address this question, we generated a series of deletions selectively targeting the three TSSs at A, B, and C followed by E/L Repli-seq to measure RT and BrU-seq to measure nascent transcription (**Fig. 4** and **Supplemental Figs. 3 - 4**). The CTSS deletion in the WT background, whose nonsignificant effect on RT was described in **Fig. 1Ciii and D**, had little effect on transcription (**Fig. 4A and F)**. We deleted BTSS in both the WT background and in the ΔCTSS background to generate ΔBTSS and ΔBTSS.CTSS, which both resulted in barely detectable BrU-seq signal from the Morc1 gene (**Figs. 4A and Supplemental Figure 2**) and a reduction in Dppa2 and Dppa4 transcription, but only a modest impact on RT, maintaining early RT (**Fig. 4Bi and ii and Supplemental Fig. 3**).

**FIGURE 4.**
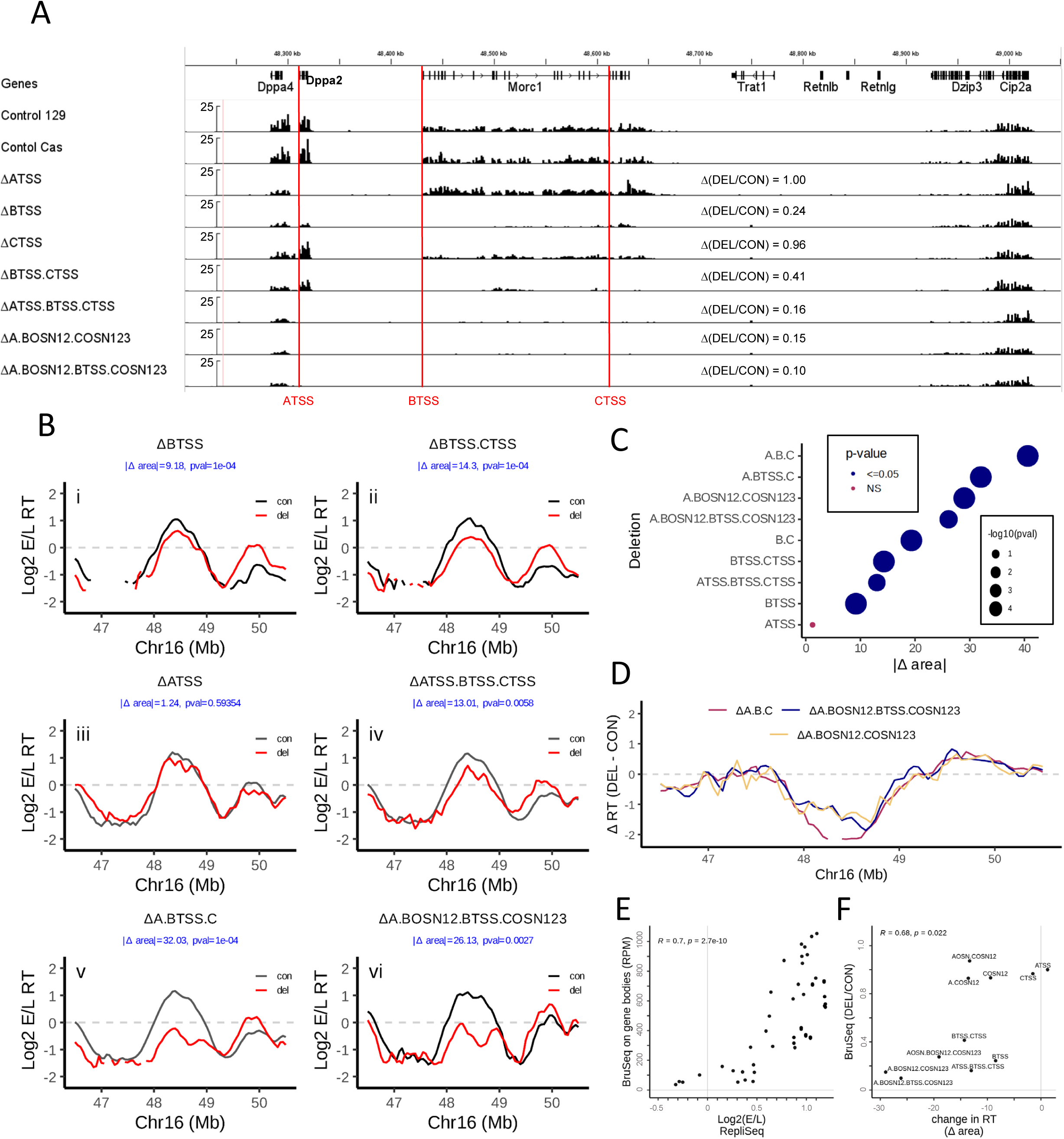
Most transcription is not required for early RT of the Dppa2/4 domain. A) Representative IGV Bru-seq tracks with TSS deletions indicated at red vertical lines. Tracks of all individual replicate BrU-seq datasets generated in this study are shown in Supplemental Fig. 5. B) RT profiles of cell lines harboring deletions (red) compared to the homologous unmodified WT allele (black) of sites within ERCE B presented as in Figure 1C. For plots in which the control (con) is grey, at least one of the clones has a deletions on both alleles and the control is an average of all control alleles in all clones in this study. C) |Δ area| between the RT profile curves for the deletions shown in B and the significance of their delay in RT compared to their corresponding control (see Materials and Methods). D) Comparison of RT changes [ΔRT (DEL-CON)] in the Dppa domain comparing ΔA.BOSN12.BTSS.COSN123 to ΔA.BOSN12.COSN123 and ΔA.B.C. E-F) Scatter plots of all clones for which both RepliSeq and BruSeq were performed, showing (E) the correlation between the average RT of Dppa domain map coordinates 48.2-48.6 Mb for each clone and the corresponding BruSeq signal over all three gene bodies (Dppa2/4 and Morc1), and (F) the correlation between the relative change in BruSeq (DEL/CON) and the change in RT (Δ area).

Next, to eliminate transcription from the Dppa2 gene, we deleted ATSS. ΔATSS alone resulted in undetectable BrU-seq signal from the Dppa2 gene and a reduction in Dppa4 with no effect on Morc1 transcription (**Figs. 4A, and Supplemental Figure 4**) and a nonsignificant effect on RT (**Fig. 4Biii and Supplemental Fig. 3**). Deletion of all three TSSs (ΔATSS.BTSS.CTSS) resulted in undetectable BrU-seq signal from Dppa2 and large reductions in BrU-seq signal from Morc1 and Dppa4 (**Figs. 4A and Supplemental Fig. 4**) with a modest change in RT similar to ΔBTSS.CTSS (**Fig. 4Biv, 4C, 4F and Supplemental Fig. 3**). Importantly, despite the nearly complete elimination of all transcription in this triple TSS deletion, Dppa domain replication is still occurring in the first half of S phase (**Fig. 4Biv)**. Thus, we conclude that, while transcription plays a role in advancing RT, most transcription is not necessary for early replication at the Dppa domain when subERCEs are intact.

Because ΔBTSS eliminates almost all transcription of the 1Mb Morc1 gene (**Fig. 4A**) and ∼76% of total BrU-seq signal in the domain, but remains early replicating (**Fig.4bi, 4C and 4F**), we generated a BTSS deletion in the ΔAC background, and found that ΔA.BTSS.C resulted in a significantly greater delay than ΔAC (**Fig. 4Bv, Supplemental Fig. 3viii vs. 3ix and Supplemental Table 4**), from slightly early-mid S to slightly late-mid S. These data support a role for BTSS in advancing RT when A and C are absent.

Recall from **Figure 3Civ** that deletion of all OSNs within all three ERCEs failed to completely delay RT equivalent to ΔA.B.C. Since the BTSS remains in this configuration and is the most consequential TSS deletion for both RT and transcription, we determined whether the BTSS was responsible for both transcription and the mid-late RT in this clone. We found BrU-seq signal was nearly eliminated in ΔABOSN12.COSN123 even though BTSS and CTSS are still present (**Fig. 4A and F and Supplemental Fig. 4**). Deletion of BTSS to create ΔA.BOSN12.BTSS.COSN123 did not significantly delay RT further (**Fig. 4Bvi, C-D and F, Supplemental Table 4**). These data demonstrate that subERCEs play a role in transcription but when transcription is already crippled by their absence, the BTSS no longer plays a role in RT, suggesting that the role of BTSS is likely through transcription.

Overall, we conclude that subERCEs have roles in both transcription and RT but that when nearly all transcription is eliminated through deletion of the TSSs, they can still maintain early replication, uncoupling these two functions of subERCEs.

### Early RT is independent of where replication initiates

We previously (Sima *et al*., 2019) aligned the ERCE-containing deletions to available small nascent strand microarray mapping data in mESCs (Cayrou *et al*, 2015) and reported that one of many origins in the Dppa domain was detected within the left half of the C deletion and none in the A or B deletions (**Supplemental Fig. 1**). Since then, several new mESC replication initiation datasets using various methods have been published, prompting us to re-examine their alignment with the more refined sites of RT ac-tivity within ERCEs. The top two data tracks in **Figure 5A** align the Dppa locus and its ERCEs with data for High resolution Repli-seq (Zhao *et al*, 2020) and Ok-seq (Petryk *et al*, 2018), which map the first sequences to replicate and the population-averaged polarity of replication, respectively, both at ∼50kb resolution. Each detect large regions of dispersed initiation activity termed “initiation zones” (IZs), although Repli-seq IZs score all earliest 50kb windows above background while the Ok-seq IZ call is a single site marking the “center of gravity” of surrounding initiation activity (Petryk *et al*., 2018). In genome-wide data, all mESC Ok-seq IZ calls are early replicating, and align with the early replicating IZs called by High resolution Repli-seq (Zhao *et al*., 2020). Indeed, at the Dppa locus, the transition in Okazaki fragment polarity (Ok-seq IZ) is near the center of the earliest DNA synthesis IZ detected by High resolution Repli-seq. In fact, the IZ called by Repli-seq (Zhao *et al*., 2020) extends from one border of strong Ok-seq polarity to the other border of opposite polarity (**Fig. 5A)**, coinciding with a broad region of reduced Okazaki fragment polarity characteristic of dispersed initiation activity (red dashed box in **Fig. 5A**), and encompassing all three ERCEs A, B and C. Together, these two very different methods provide high confidence that initiation at the Dppa locus is distributed throughout the entire region encompassed by the ERCEs.

**FIGURE 5.**
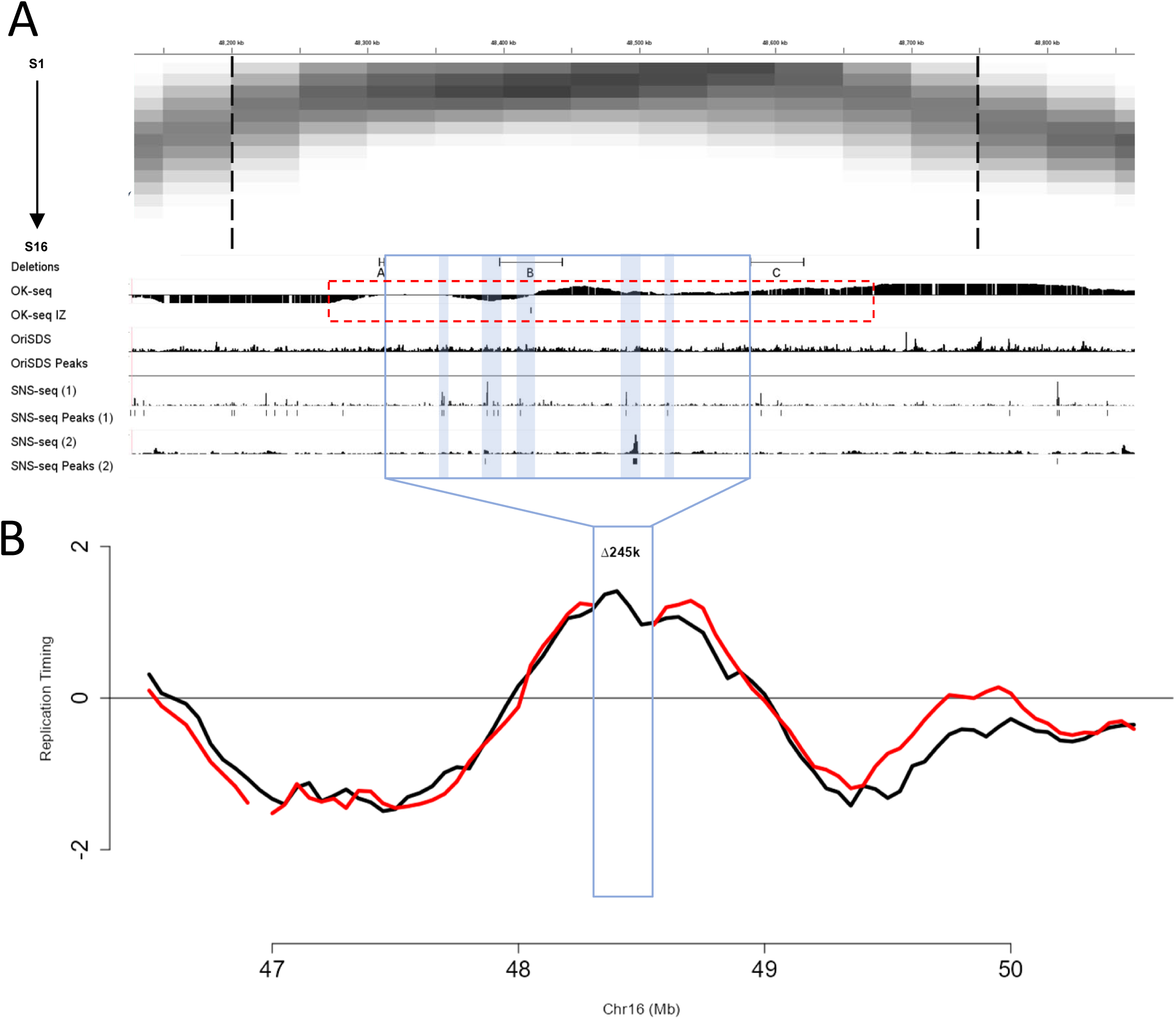
ERCEs and replication origins. A) IGV tracks showing: (Top) a heat map of read coverage in 50kb genomic windows from high-resolution Repli-seq of the musculus allele in F121-9 mESCs, with the IZ call indicated by the vertical black dashed lines, which identifies the locally earliest 50kb bins with read coverage significantly above background, (Zhao et al., 2020). (Middle) Deletions and signal tracks followed by peak calls for Okazaki fragment sequencing (Ok-seq) data (Petryk et al., 2018). The “IZ Peak” calls in Ok-seq data represent the point where the population average of fork directions switches polarity. The red dashed box represents the region of reduced uni-directional replication (increased bi-directional replication), indicative of origin firing. (Bottom) Signal tracks followed by peak calls for three small nascent strand sequencing (SNS-seq) datasets. OriSDS is a revised SNS method in which only bi-directional small nascent strands are scored as peaks (Pratto et al., 2021). SNS dataset (1) and (2) are from (Prorok et al., 2019) and (Jodkowska et al., 2022), respectively. B) Log2(E/L) Repli-seq of a clone with a 245kb deletion encompassing most of the efficient SNS origin peak calls (Sima et al., 2019). Blue box extending up into (A) expands the deleted region to show positions of SNS and Ok-seq IZ peak.

Also shown in **Figure 5A** are three recently published mESC small nascent strand sequencing (SNS-seq) mapping datasets (Jodkowska *et al*, 2022; Pratto *et al*, 2021; Prorok *et al*, 2019), which map initiation sites to kilobase resolution. One of these datasets (SNS-seq 1) detects peaks throughout the IZ, including two peak calls within the left half of C and one in the left side of B. The other two datasets (OriSDS and SNS-seq 2) detect 0 and 2 peaks in the entire Dppa domain, none of which reside within the original large ERCE-containing deletions (**Fig. 5A**). Importantly, none of the SNS-seq datasets detected peaks within 10kb of any of the subERCEs. Together with the Repli-seq and Ok-seq data, the most parsimonious conclusion is that origin activity in the Dppa locus is distributed throughout the domain, with some sites firing at efficiencies above the threshold of SNS-seq peak detection. To evaluate the importance of efficient sites of initiation for Dppa domain RT, we aligned the position of a large (∼245kb) deletion made in our previous work (Sima *et al*., 2019), that had no detectable effect on RT (**Fig. 5B**). This deletion removes all DNA between the A and C elements, encompassing the majority of detectable SNS activity throughout the Dppa domain. Thus, initiation frequencies must necessarily be re-distributed in this deletion to the remaining flanking regions; this suggests RT is regulated independently of the choice of sites used to initiate replication.

## DISCUSSION

*Cis*-acting elements controlling large-scale (> hundreds of kb) chromosome structure and function have proven elusive. We recently identified ERCEs as *cis*-elements necessary for early RT, and found that the same chromosome segments harboring ERCEs were also necessary for domain-confined transcription, 3D architecture and A/B compartmentalization. Most of our deletions were over 30kb, making it difficult to determine whether early replication activity could be uncoupled from these other activities. Here we have performed a series of fine deletions within the 3 ERCES residing in the ∼400kb Dppa2/4 gene-containing domain. We demonstrate that a set of 6 relatively small (140-3483 bp; 7384 kb total) deletions can mostly recapitulate the very early to very late replication timing switch observed with the larger deletion series (which deleted 88.7 kb total). These 6 deletions contain 7 sites, we term subERCEs, which are bound by diverse master TFs. Although the 3 subERCEs in ERCE C account for its RT activity in most deletion series, deletion of all 7 subERCEs does not render the entire domain homogenously very late replicating, such as what we see with ΔA.B.C. At present we do not know what alternative activity is contributing to the remaining late-middle RT in these clones. Deletion of all subERCEs eliminates almost all transcription in the locus, despite retaining all domain TSSs. On the other hand, deleting three TSSs (ΔATSS.BTSS.CTSS), leaving 6/7 subERCEs intact (the seventh is ATSS, which may also be a subERCE), also eliminates almost all transcription, but the Dppa locus replicates early in this clone. Thus, subERCEs have roles in both transcription and RT but, when nearly all transcription is eliminated through deletion of the TSSs, they can still maintain early replication, uncoupling these two functions of subERCEs. Note that a 140bp deletion of only the BTSS substantially reduces transcription of the long Morc1 gene, with a significant effect on RT. However, when transcription was already reduced by deleting all 7 subERCEs, further deletion of the BTSS had no additional effect on RT, suggesting that the role of BTSS in RT is strictly through transcription, distinguishing it from subERCEs. Together, we conclude that subERCEs are compound elements consisting of multiple independent sites that function to maintain early RT through roles in both stimulating transcription and separately by as yet undefined mechanisms that can function independent of transcription. Thus, changes in transcription factors during cell fate changes can alter RT without affecting transcription and may cooperate with transcription to elicit robust changes in RT.

### ERCEs consist of multiple smaller “subERCEs”

Our original deletion study (Sima *et al*., 2019) found that very early replication requires at least two ERCEs. Here, we show here that each of the ERCEs consists of multiple smaller elements, identified by OSN co-binding. However, it is clear that the OSN TFs themselves are not sufficient for ERCE RT activity, since two additional strong OSN sites (Y and Z) remain intact in all of our late S phase replicating deletion clones. Since the term OSN is thus misleading with respect to RT activity, we have introduced the term “subERCEs” for elements within ERCEs that contribute to the RT activity of ERCEs independent of the transcriptional contribution of ERCEs to RT (i.e. BTSS). Importantly, both motif analysis and ChIP data identify a large number of TFs that bind to subERCEs (**Supplemental Figure 5A-C**). In fact, it has been shown that the diversity of TF binding sites is more important for predicting enhancer activity than the specific identity of TFs (Singh *et al*, 2021). However, X, Y and Z and even CTSS also contain many TF binding sites (**Supplemental Figure B**), so there is no obvious combination of TFs that stands out as distinguishing subERCEs and the number of elements in our study is too small to find a TF signature unique to subERCEs. **Supplementary** Figure 5D shows recently published high-resolution H3K27ac HiChIP and Micro-C data for the Dppa locus in mESCs (Jusuf *et al*, 2025; Kraft *et al*, 2022). Consistent with our previous capture Hi-C data (Sima *et al*., 2019), interactions are seen between ERCEs A and C and between B and C, but not between A and B. Consistent with the activity of subERCEs being linked to their interactions, interactions are enriched at the subERCE sites within the ERCEs, albeit in all cases TSSs are in close proximity to the subERCEs. Interestingly, Y and Z show no significant interactions with any subERCE in either the

HiChIP or micro-C data, and X only has one contact with ERCE A. Thus, it is possible that inter-ERCE interaction distinguishes subERCEs. It should also be noted that a recent study demonstrated that superenhancers contain bioinformatically equivalent but functionally distinct “facilitators” that potentiate enhancer activity dependent upon their positions within the superenhancer (Blayney *et al*, 2023), raising intriguing parallels to subERCEs within ERCEs. Facilitators were identified using a whole-locus synthetic biology-enabled engineering strategy designed to overcome the difficulties of introducing multiple independent mutations in a single allele. Our conventional CRISPR deletion approach has refined the size of ERCEs and identified a complexity of cooperative elements akin to super-enhancers. However, this approach is laborious and precisely positioning Cas9 PAM/gRNA sequences is not possible. In short, our work described here has revealed the complex nature of ERCEs. Novel approaches will be needed to elucidate mechanisms by which subERCEs maintain early replication, potentially coordinating changes in RT with transcription during differentiation.

### ERCEs and transcription

We demonstrate that subERCE RT activity can maintain early replication independent of its roles in transcription, uncoupling these two identified activities of ERCEs. Even after all ERCE-linked transcriptional activity (all the transcription eliminated in ΔA.B.C) and 90% of total transcriptional activity throughout the Dppa domain, is eliminated by deleting ERCE-resident TSSs, the domain remains early replicating. However, our results also suggest a role for transcription in the robust maintenance of Dppa domain RT. It has been shown that high levels of transcription, or transcription of long genes can be sufficient but not necessary to drive early RT during differentiation (Vouzas *et al*, 2025), suggesting something other than transcription, such as ERCEs, function when transcription is not activated. Moreover, transcription of small, lowly transcribed genes such as Dppa2/4 may not influence RT at all (Blin *et al*, 2019; Vouzas *et al*., 2025). Our work shows that, at the Dppa locus, the BTSS deletion, that eliminates most domain transcription including the long Morc1 gene, causes a significant delay in RT, but is not enough to erase early replication, even after additional deletion of ATSS and CTSS, so it contributes to how early the Dppa locus is, but is not necessary for early replication. Thus transcription is one of at least two independent mechanisms maintaining Dppa RT. The effect of transcription on RT may be direct (Vouzas *et al*., 2025). Alternatively, transcription could result in alterations in the positions of the replicative MCM helicase complexes that are known to be cleared from the bodies of active genes (Chen *et al*, 2019; Hu & Stillman, 2023; Lichauco *et al*, 2025; Liu *et al*, 2021; Sasaki *et al*, 2006; Scherr *et al*, 2022). Thus, silencing transcription could result in a broad distribution of origin firing across the large Morc1 gene body, each of which is equally early firing but used infrequently, so that any given genomic bin appears to replicate later in ensemble data. Indeed, this model could explain why the 245k deletion that removes most of the Morc1 gene has no effect on RT (**Figure 5**), while the 140-bp BTSS deletion significantly delays RT (**Figure 3**); the 245k deletion removes most of the Morc1 gene body, which may re-focus MCM and initiation to a smaller genomic region. Regardless of the mechanism, the most parsimonious interpretation of our results is that promoters, such as BTSS, advance RT strictly by driving transcription, while subERCEs advance RT both through their role in transcription, likely functioning as enhancers or facilitators, and through a transcription-independent mechanism. Until we understand that mechanism, it will remain difficult to bioinformatically identify subERCEs.

### ERCEs and replication origins

Although replication initiates at sites dispersed throughout the Dppa locus, RT is not linked to any specific sites of initiation. This is consistent with a body of literature uncoupling RT from origin specification, discussed in depth elsewhere (Rivera-Mulia & Gilbert, 2016). Indeed, the role of any site-specific origin of replication in chromosome biology has been difficult to substantiate. There is evidence that altered origin usage can influence the frequency of trinucleotide repeat expansion by altering the polarity of replication forks passing through the repeats (Gerhardt *et al*, 2014). There is also evidence of an early to late RT change upon deletion of a strong SNS-seq peak in mouse B cells (Malzl *et al*, 2023), although ReMap2022 ChIP data reveals that the deleted site binds diverse TFs, including B cell master regulators such as Pax5, and so could contain one or more subERCEs. Here, we demonstrate that we can delete the most efficient origins within an initiation zone, leaving two ERCEs behind, with no detectable effect on RT.

To the non-expert, the discordance between origin peak calls in **Figure 5** may seem surprising. In fact, it is typical of most genomic regions in mammalian cells, particularly developmentally regulated domains that share properties with late domains, including dispersed origin distribution (Besnard *et al*, 2012; Dileep *et al*, 2015; Takebayashi *et al*., 2012b), likely due to a combination of technical noise and the biological background of dispersed initiation. The usage of sites of replication initiation in mammalian cells is highly stochastic (Bechhoefer & Rhind, 2012; Carrington *et al*, 2024; Hyrien, 2015; Wang *et al*, 2021). Single molecule methods show that 20% of origins are organized in clusters averaging 45kb termed Initiation Zones (IZs), within which replication initiates at one or a few of many more potential sites. The remaining 80% of initiations are scattered throughout the genome outside of the IZs and fire at an undetectably low frequency. This has made origin mapping very difficult, as single molecule methods are challenging and population-based methods that achieve kilobase resolution such as SNS-seq (Jodkowska *et al*., 2022; Pratto *et al*., 2021; Prorok *et al*., 2019) and initiation site sequencing (Ini-seq; (Guilbaud *et al*, 2022) detect only those with sufficient efficiency to give a signal above noise. There are, however, some exceptionally efficient and reproducibly detectable “core origin” sites (Akerman *et al*, 2020), which mainly reside within gene-rich domains replicated very early across cell types. By contrast, lower resolution methods Ok-seq (Petryk *et al*, 2016) and High resolution Repli-seq (Zhao *et al*., 2020), which detect sites where many origins are clustered into IZs, give high concordance between replicates and even between these two very different data types (**Figure 5** and (Zhao *et al*., 2020). In summary, IZs are broad regions of chromatin where replication initiation is enriched and the entire Dppa replication domain can be considered one (or possibly two) large IZ. ERCEs, on the other hand, are discrete elements required for large-scale domain structure and function. Taken together, we favor a model in which ERCEs establish a subnuclear environment that promotes initiation stochastically at available sites within their realm of influence, the replication domain (**Figure 6**; (Dimitrova & Gilbert, 1999; Gilbert, 2001; Sima *et al*., 2019).

**FIGURE 6.**
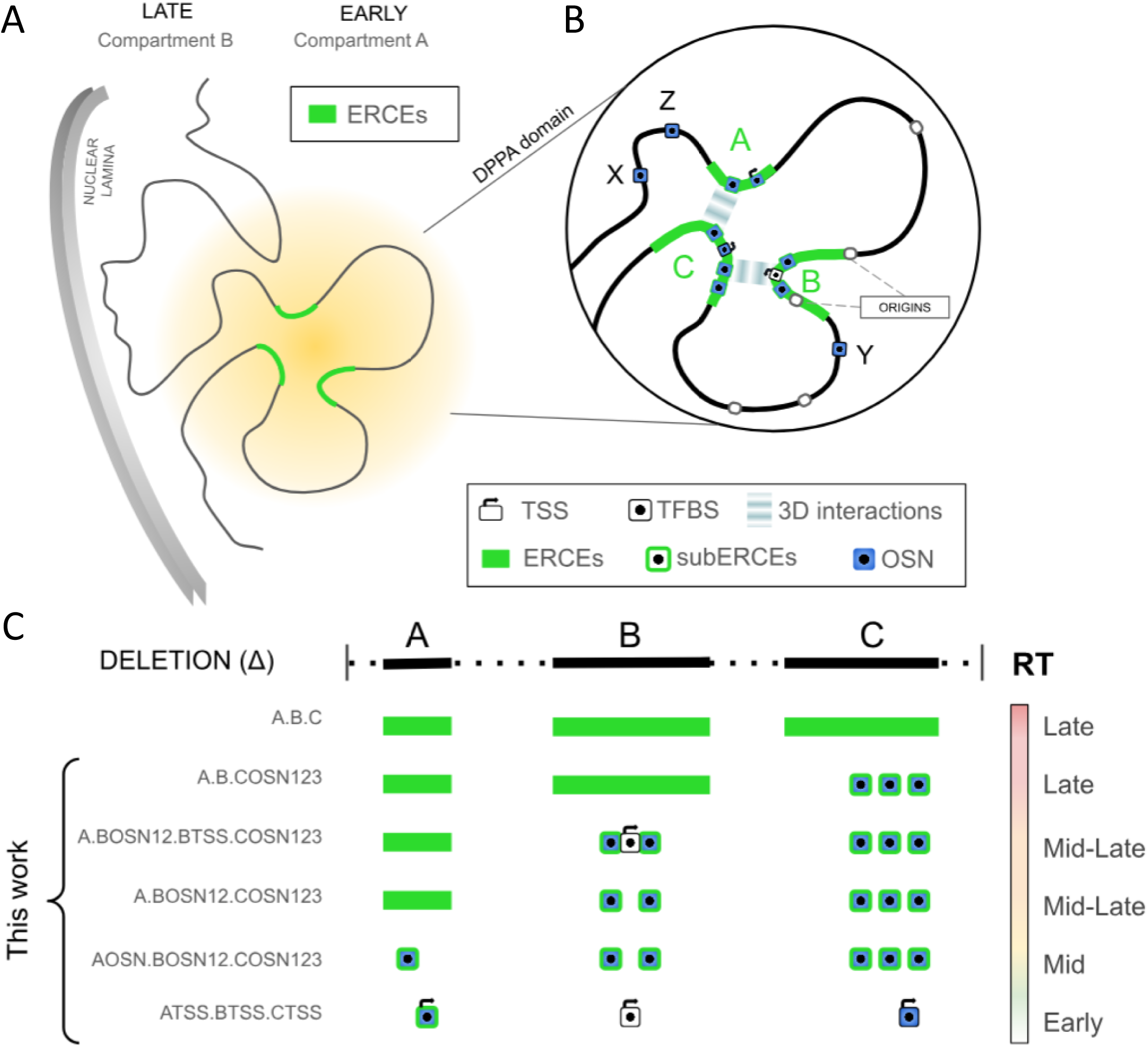
Uncoupling activities within ERCEs. Working model for ERCE function, illustrating the various activities of candidate regulatory elements subject to deletion analysis in this work, while retaining the basic tenets of our original proposed model (Sima et al., 2019). A) Original version of the model adapted from Sima et. al., 2019, showing a hypothetical structure of the Dppa replication domain, in which interactions between ERCEs seed the assembly of a microenvironment (yellow halo) that results in interactions with the A chromatin compartment and increases the local probability of origin firing. B) Revised model showing the positions and interaction of all dissected elements in this study (marked by boxes) and classified according to function. Bubbles represent stochastically used replication origins within an initiation zone that aligns with ERCE B. Interactions occur between subERCEs A-C and B-C but are rarely detected between A-B. C) Schematic representation of relevant compound deletions and their resulting effects on RT, highlighting the contribution of this work towards a detailed understanding of ERCE function.

### Co-opting TFs as a means to regulate RT during development

Our results strongly implicate master transcription factor binding sites in developmental control of RT. Coupled transcription and RT activities of a subset of enhancers suggest one possible reason why RT and transcriptional regulation have been so hard to untangle, and a partial explanation for why there are so many more cell type specific TF binding sites than there are essential transcriptional enhancers (Lo *et al*, 2022; Rothenberg, 2022). Our results also suggest that cell type specific ERCEs could respond to changes in transcriptional regulatory networks during differentiation to trigger replication timing changes, thus remodeling the epigenomic composition of their resident domain to create an environment more conducive for transcription (Klein *et al*., 2021). Our results are also consistent with a role for transcription in advancing RT, as reported by others (Blin *et al*., 2019; Therizols *et al*, 2014), and suggest that these mechanisms can act independently. Potentially then, either transcription or ERCEs could initiate RT advances during cell fate specification, while the alternative mechanism could serve as a positive feedback to drive stable transitions in epigenetic state in either temporal order. Testing this model in mammalian cells will require further understanding of mechanisms by which ERCEs function.

Figure 6 illustrates a speculative model for ERCE function, modified from its original version (Figure 6A; (Sima *et al*., 2019)) to include the discovery of TF-rich subERCEs, as well as recently published origin mapping and Micro-C/HiChIP data (Figure 6B). Figure 6C summarizes key observations in this report. Here we demonstrate that ERCEs consist of multiple subERCEs, which are sites of diverse cell-type specific TF binding. The subERCEs account for most of the RT advancing activity of ERCEs. ERCEs were originally discovered by virtue of their CTCF-independent interactions (Sima *et al*., 2019). We show here that ERCE interactions are focused on the more localized subERCEs (**Suplementary** Figure 5D), while Y and Z interact do not interact with subERCEs. Interestingly, synthetic sequences that can advance RT in chicken DT40 cells, when inserted at distant sites, have been shown to interact in 3D space and when doing so they synergize in promoting early replication (Brossas *et al*, 2020). We propose that subERCEs create a sub-nuclear microenvironment, potentially through (micro)phase separation (Goel *et al*, 2024), that promotes initiation of replication within the structurally organized replication domain. This model for subERCE activity is purposely reminiscent of binding sites for the Fkh1/2 TFs in budding yeast, which reside near a set of early firing origins and promote early RT by dimerization-mediated interactions, independently of their role in transcription (Ostrow *et al*., 2017). In fact, Oct4 and Nanog, as well as Yamanaka factor Klf4, have been shown to mediate chromatin looping (Choi *et al*, 2022; de Wit *et al*, 2013; Di Giammartino *et al*, 2019), and several studies have shown that master TFs form 3D hubs to regulate cell identity (Hu *et al*, 2023; Liu *et al*, 2023; Madsen *et al*, 2020; Wang *et al*, 2022a; Wang *et al*, 2022b; Winick-Ng *et al*, 2021). FoxP1, a mammalian TF that possesses the same dimerization motif as Fkh1/2 (Ostrow *et al*., 2017), is highly expressed in ESCs and required for pluripotency (Gabut *et al*, 2011). Moreover, both Sox2 and Klf5 interact with the pluripotency-specific isoform of FoxP1 (Göös *et al*, 2022), which contains a self-dimerization motif homologous to that of budding yeast Fkh1/2. Thus, mechanisms regulating constitutive replication timing in budding yeast may inform us as to how RT changes occur during mammalian cell fate transitions. It is now of high interest to determine whether one or more of the TFs bound to subERCEs mediates interactions between ERCEs, and if so, whether these interactions regulate RT independently of their role in transcription.

## MATERIALS AND METHODS

### Publicly-Available Datasets Used

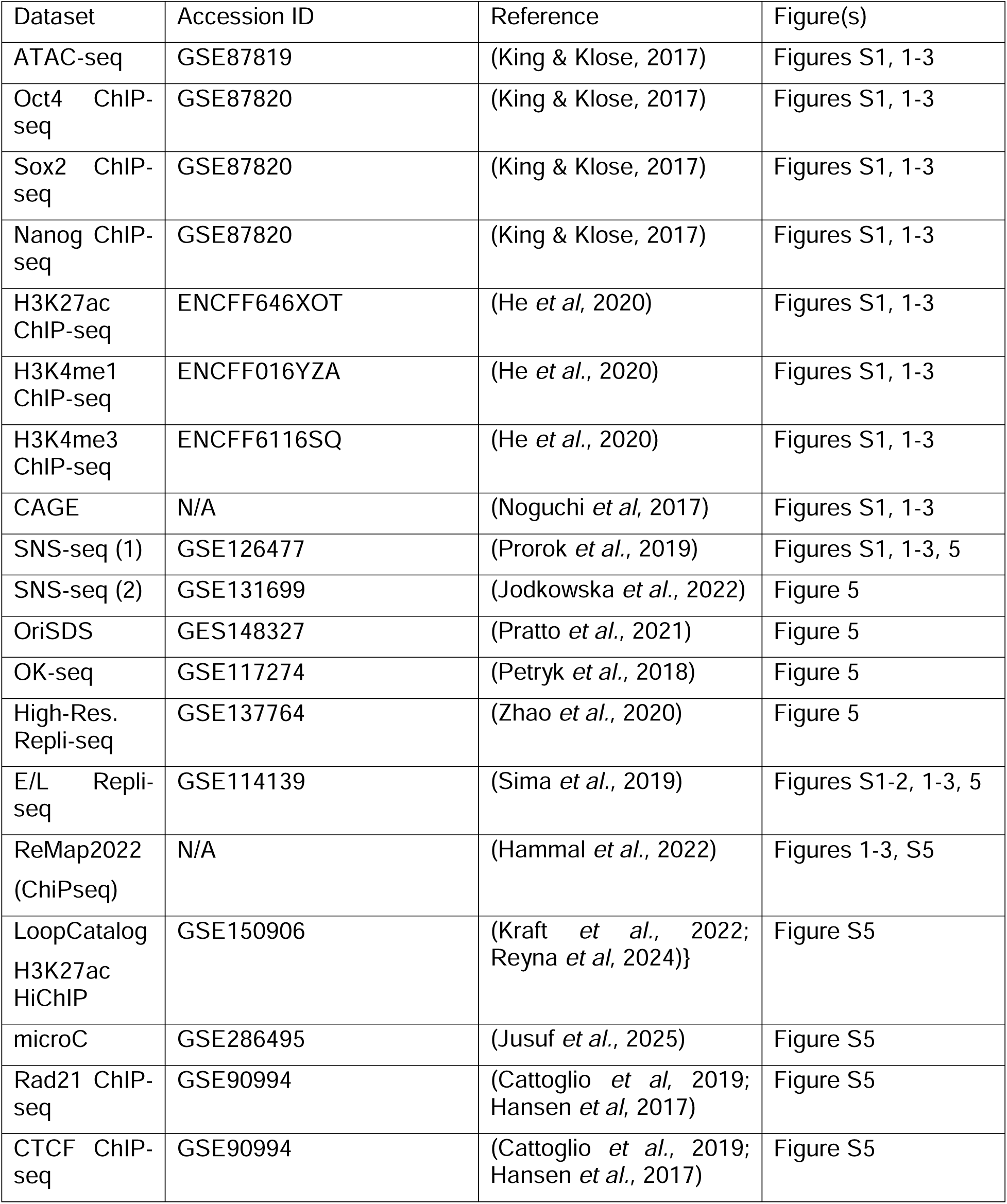

| Dataset | Accession ID | Reference | Figure(s) |
| --- | --- | --- | --- |
| ATAC-seq | GSE87819 | (King & Klose, 2017) | Figures S1, 1-3 |
| Oct4 ChIP-seq | GSE87820 | (King & Klose, 2017) | Figures S1, 1-3 |
| Sox2 ChIP-seq | GSE87820 | (King & Klose, 2017) | Figures S1, 1-3 |
| Nanog ChIP-seq | GSE87820 | (King & Klose, 2017) | Figures S1, 1-3 |
| H3K27ac ChIP-seq | ENCFF646XOT | (He *et al.*, 2020) | Figures S1, 1-3 |
| H3K4me1 ChIP-seq | ENCFF016YZA | (He *et al.*, 2020) | Figures S1, 1-3 |
| H3K4me3 ChIP-seq | ENCFF6116SQ | (He *et al.*, 2020) | Figures S1, 1-3 |
| CAGE | N/A | (Noguchi *et al.*, 2017) | Figures S1, 1-3 |
| SNS-seq (1) | GSE126477 | (Prorok *et al.*, 2019) | Figures S1, 1-3, 5 |
| SNS-seq (2) | GSE131699 | (Jodkowska *et al.*, 2022) | Figure 5 |
| OriSDS | GES148327 | (Pratto *et al.*, 2021) | Figure 5 |
| OK-seq | GSE117274 | (Petryk *et al.*, 2018) | Figure 5 |
| High-Res. Repli-seq | GSE137764 | (Zhao *et al.*, 2020) | Figure 5 |
| E/L Repli-seq | GSE114139 | (Sima *et al.*, 2019) | Figures S1-2, 1-3, 5 |
| ReMap2022 (ChIPseq) | N/A | (Hammal *et al.*, 2022) | Figures 1-3, S5 |
| LoopCatalog H3K27ac HiChIP | GSE150906 | (Kraft *et al.*, 2022; Reyna *et al.*, 2024)} | Figure S5 |
| microC | GSE286495 | (Jusuf *et al.*, 2025) | Figure S5 |
| Rad21 ChIP-seq | GSE90994 | (Cattoglio *et al.*, 2019; Hansen *et al.*, 2017) | Figure S5 |
| CTCF ChIP-seq | GSE90994 | (Cattoglio *et al.*, 2019; Hansen *et al.*, 2017) | Figure S5 |

### Cell Culture

mESCs were cultured, maintained, and passaged as previously described (Sima *et al*., 2019). Briefly, cells were cultured on 0.1% gelatin-coated dishes in serum-free 2i/LIF media.

### Cell Line Engineering and Genotyping

Cells were transfected using P3 Primary Cell 4D Nucleofector X Kit (Lonza, V4XP3024) or XFect Transfection Reagent (Takara, 631318) with two px458 (Addgene, 48138) plasmids containing gRNA sequences targeting the deletion breakpoints. Three days after transfection, GFP-positive cells were single-cell sorted by FACS into 96-well plates. Ten to fourteen days after sorting, colonies were picked and expanded into duplicate 96well plates and PCR screened. Primers upstream and downstream of the breakpoints are used to identify deletion mutants, and for large (>2kb) deletions, primers within the deleted sequences are used to identify heterozygous mutants. Deletion PCR fragments were then sanger-sequenced to determine the haplotype of the deletion allele which was then verified by NGS. The sgRNA and PCR primer information can be found in **Supplemental Table 1**. Deletion coordinates and sgRNAs used for each deletion can be found in **Supplemental Table 2**. Individual cell line genotypes as determined by Sanger sequencing and NGS analysis can be found in **Supplemental Table 3**. Cell line authentication was done by analyzing the whole genome repli-seq data and all lines are free of mycoplasma.

### Repli-seq Library Preparation and Data Analysis

Repli-seq was performed as previously described (Marchal et al. 2018). Briefly, cells were labeled with 100uM BrdU (Sigma Aldrich, B5002) for 2 hours and then fixed in 75% ethanol. Fixed cells were then sorted by FACS into early-S and late-S fractions based on propidium iodide staining of DNA. DNA was then purified from each fraction, 46 sheared using Covaris ME220, and used to construct Illumina sequencing libraries with NEBNext Ultra Library DNA Prep Kit for Illumina (NEB, E7370). BrdU-labeled nascent DNA library fragments were then enriched by immunoprecipitation with anti-BrdU antibody (BD, 555627). IP products are then PCR amplified and indexed. A minimum of 30 millions reads/read pairs per library were requested. Raw sequencing data were first quality and adaptor trimmed using Trim Galore (https://github.com/FelixKrueger/TrimGalore) (Martin, 2011) and then aligned to the reference genome with bowtie2 (Langmead & Salzberg, 2012). Reads were aligned to a custom N-masked mm10 genome for subsequent haplotype phasing with SNPsplit (Krueger & Andrews, 2016). Aligned reads were then quality filtered and duplicate reads removed using Samtools (Li *et al*, 2009). Processed aligned reads were then counted in 50kb windows across the entire genome for both early- and late-fraction using bedtools (Quinlan & Hall, 2010) and then a log2 ratio of early-to-late read counts for each window was calculated to generate raw timing files. Raw E/L data was then scaled with R, quantile normalized with R package preprocessCore (Bolstad, 2023) and loess-smoothed in 300kb windows to generate the final replication timing profiles used for plotting and analysis. All individual datasets are plotted in their respective groups in **Supplemental** Fig. 3.

### Statistical Analysis of RT changes

To quantitatively measure the effect of a deletion in delaying the replication timing of the DPPA domain, we implemented a randomization-based statistical approach. We simply ask the question of whether the delay in RT (computed as the area between curves representing control and deletion alleles) observed from a given real deletion is significantly larger than expected by chance. First, we calculated the area under the averaged Δ RT curve (deletion control) for each target deletion in the DPPA domain (chr16 47.5Mb → 49.5Mb) using the AUC function in R. For the clones without a WT internal control (homozygous deletions), we averaged the RT profiles from all available WT alleles from all the clones in this work to generate the aggregate WT curve. To estimate random fluctuations in the RT data between the two alleles, we built a background model by randomly sampling the genome to obtain ten thousand regions size-matched to the query (2Mb DPPA domain in this case), excluding previously characterized genomic regions with allele-specific RT in the 129/cas hybrid line (Rivera-Mulia, et al. Genome Research 2018). We utilized a p-value cutoff of 0.05 to determine which deletions significantly delay the RT of the DPPA domain. Similarly, to compare between two deletions, we employed this approach to test whether the change in RT of two distinct deletions is statistically equivalent, by comparing the ΔΔRT = ΔRT1ΔRT2 with the same randomization-based framework as described above. We developed a generic code that can assess the significance of RT change for any query region with options to allow choosing sampling size, statistical test, and consider blacklisted regions. We made these scripts available on Github: https://github.com/ay-lab/SignifRT. The statistics for all datasets and a graphical explanation of the method used can be found in **Supplemental** Figure 2. All pairwise statistical comparisons can be found in **Supplemental Table 4**.

### Bru-seq Library Preparation and Data Analysis

Bru-seq was performed as previously described (Bedi *et al*, 2020; Paulsen *et al*, 2014). Briefly, cells were labeled with 2mM Bru (Sigma Aldrich, 850187) for 30 min and then total RNA collected using Direct-Zol RNA Miniprep Plus Kit (Zymo, R2070). Bru-labeled nascent RNA was then enriched 47 using anti-BrdU antibodies conjugated to magnetic beads. Nascent RNA was then fragmented at 85C for 10 min and converted to cDNA using NEBNext Ultra II RNA First Strand Synthesis Module (NEB, E7771) and NEBNext Ultra II Directional Second Strand Synthesis Kit (NEB, E7550) according to manufacturer’s instructions. cDNA fragments were then constructed into libraries using NEBNext Ultra II DNA Library Prep Kit for Illumina (NEB, E7645). 100M reads/read pairs per library were requested. Raw sequencing data were processed and aligned like in E/L repli-seq, except duplicate reads are retained (Paulsen *et al*, 2014). RPM-normalized basepair coverage tracks were generated from the processed alignment files with bedtools and visualized in bigwig file format using IGV. All Bru-seq performed and analyzed in this study can be found in **Supplemental** Figure 4.

### Bru-seq Quantification and Bru-seq vs RT Analysis

RPM-normalized BruSeq signal at 1kb resolution was averaged across gene bodies of Morc1, Dppa2, and Dppa4 for each available clone. Data available in **Supplemental Table 5**. To compare against RT, the log2(E/L) of the DPPA replication domain was averaged across replicates of each genetically distinct clone for which both RepliSeq and BruSeq were performed. The change in BruSeq was computed as the fraction of signal observed in all deletion alleles over the average signal of all available control alleles for the corresponding genome. ΔRT was calculate as described above. Pearson correlations were computed using the stat_cor() function in R.

### TFBS Analyses

A bedfile containing all of the peak calls from the ReMap2022 database (https://remap2022.univ-amu.fr/) for mouse samples was downloaded and filtered for the “mESC” biotype, and all other peak calls were removed. We then aligned the peak calls for individual transcription factors (Oct4, Sox2, Nanog in Figures 1-3; all factors for Figure S5) to the genomic coordinates of our deletion sites and other sites of activity within the Dppa domain. We only counted one peak call from one dataset per site, and as such we only report the number of datasets that have at least one peak call in a given site. Motif search analysis on the deleted sequences was performed with FIMO (Grant *et al*, 2011) against the HOCOMOCOv11 Mouse Motif database (Kulakovskiy *et al*, 2018) with 1% FDR threshold.

## Supporting information

Supplementary Table 1 - gRNA and primer sequences

Supplementary Table 2 - deletion coordinates

Supplementary Table 3 - cell line genotypes

Supplementary Table 4 - pairwise comparisons

Supplementary Table 5 - Bruseq

## Acknowledgments

We thank B. Washburn and S. Miller from the Florida State University Department of Biological Sciences Molecular Core Facility for helpful advice, cloning help, and sanger sequencing services. We also thank Cindy Vied and Yanming Yang from the Florida State University College of Medicine Translational Science Lab for high-throughput sequencing services. This work was supported by NIH grants F31-AG066481 to JLT, R01-GM083337 to DMG and R35-GM128938 to FA. LHG was supported by an NIH training grant to UCSD T32-GM139790.

## Author Contributions

Conceptualization & Study Design, J.L.T & D.M.G, L.H.G & F.A.; CRISPR Cloning, J.L.T & C.A.F; Cell Culture, J.L.T, M.S.S, & T.S; Cell Line Engineering, J.L.T & M.S.S; PCR Screening, J.L.T; Repli-seq & Analysis, J.L.T, L.H.G & T.S; Bru-seq & Analysis, J.L.T, A.V, L.H.G & A.N.B; Flow Cytometry, J.L.T & K.E.A; Bioinformatics Analyses, J.L.T, L.H.G, A.C, & F.A; Wrote manuscript, J.L.T, D.M.G, L.H.G & F.A.

## SUPPLEMENTAL FIGURE LEGENDS

**Supplemental Figure 1.**
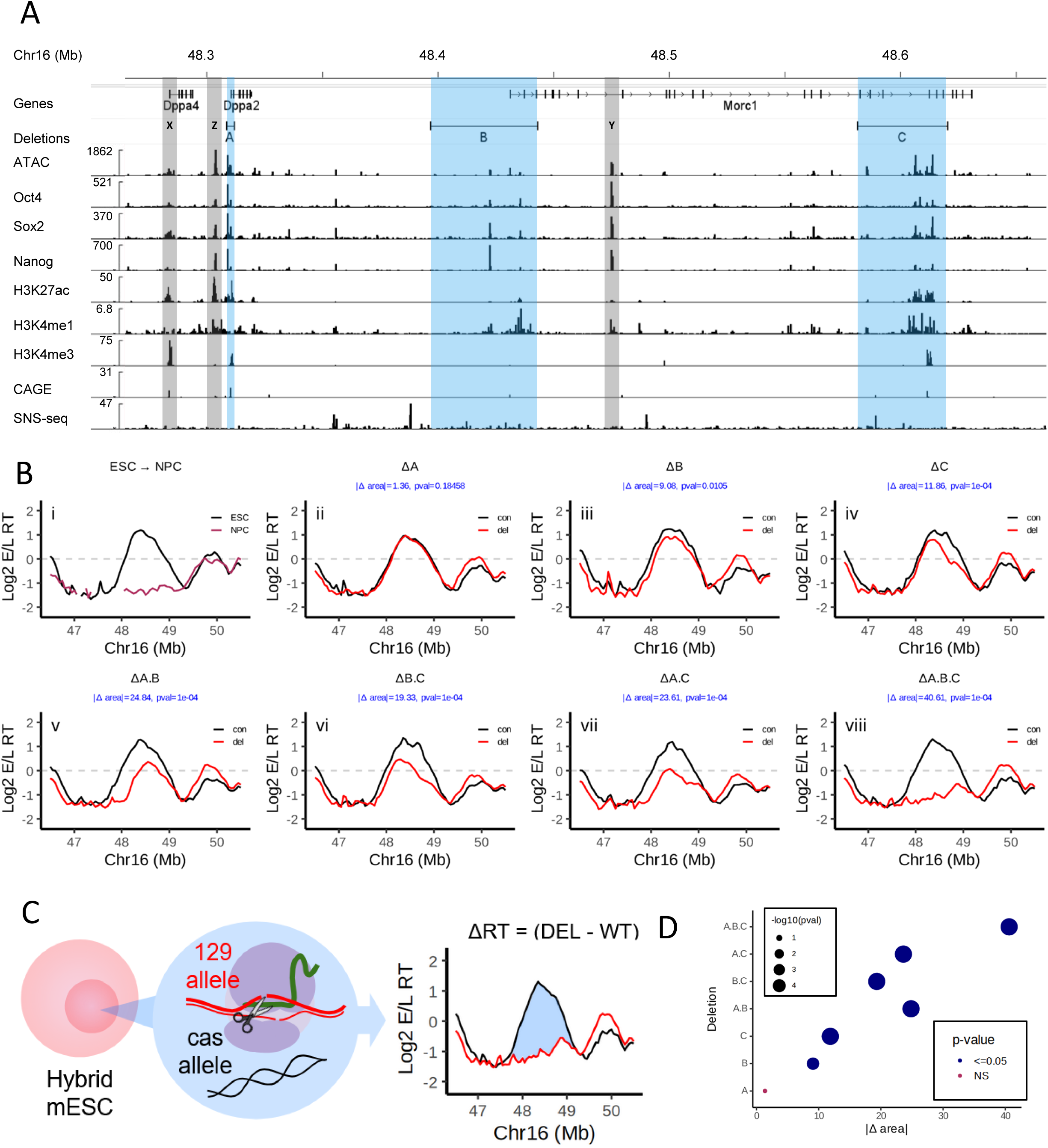
Epigenetic signature of the Dppa2/4 replication domain ERCEs. A) IGV browser tracks showing chromatin features of the Dppa2/4 domain (see **Materials and Methods** for sources). ERCEs, defined by previous deletions (Sima *et al*., 2019), are highlighted in blue. X and Y (grey highlights) represent regions that display epigenetic features of ERCEs but did not display ERCE activity in deletion analyses (Sima *et al*., 2019). B) Log2(E/L) Repli-seq (Marchal *et al*, 2018) from (Sima *et al*., 2019). The first panel (i) shows RT before and after mESC (black) to neural precursor cell (NPCs; maroon) differentiation. The remaining panels (ii-viii) show RT profiles from averaged replicates of independent CRISPR clones harboring deletions of one or more ERCEs on one allele (DEL; red) vs. the homologous unmodified allele (CON; black). Individual replicate experiments are shown in **Supplemental Fig. 3**, and the approach to assess the significance of differences in RT between a deletion allele and the corresponding WT allele are shown in **Supplemental Figure 2**. C) Schematic representation of hybrid mESC model, where one allele contains a deletion and the other serves as control, allowing us to study changes in RT by estimating the area under the ΔRT curve in the DPPA domain (See Materials and Methods). D) |Δ area| between the averaged RT profile curves of at least two replicates (shown in C) and the significance of their delay in RT compared to the corresponding WT alleles.

**Supplemental Figure 2.**
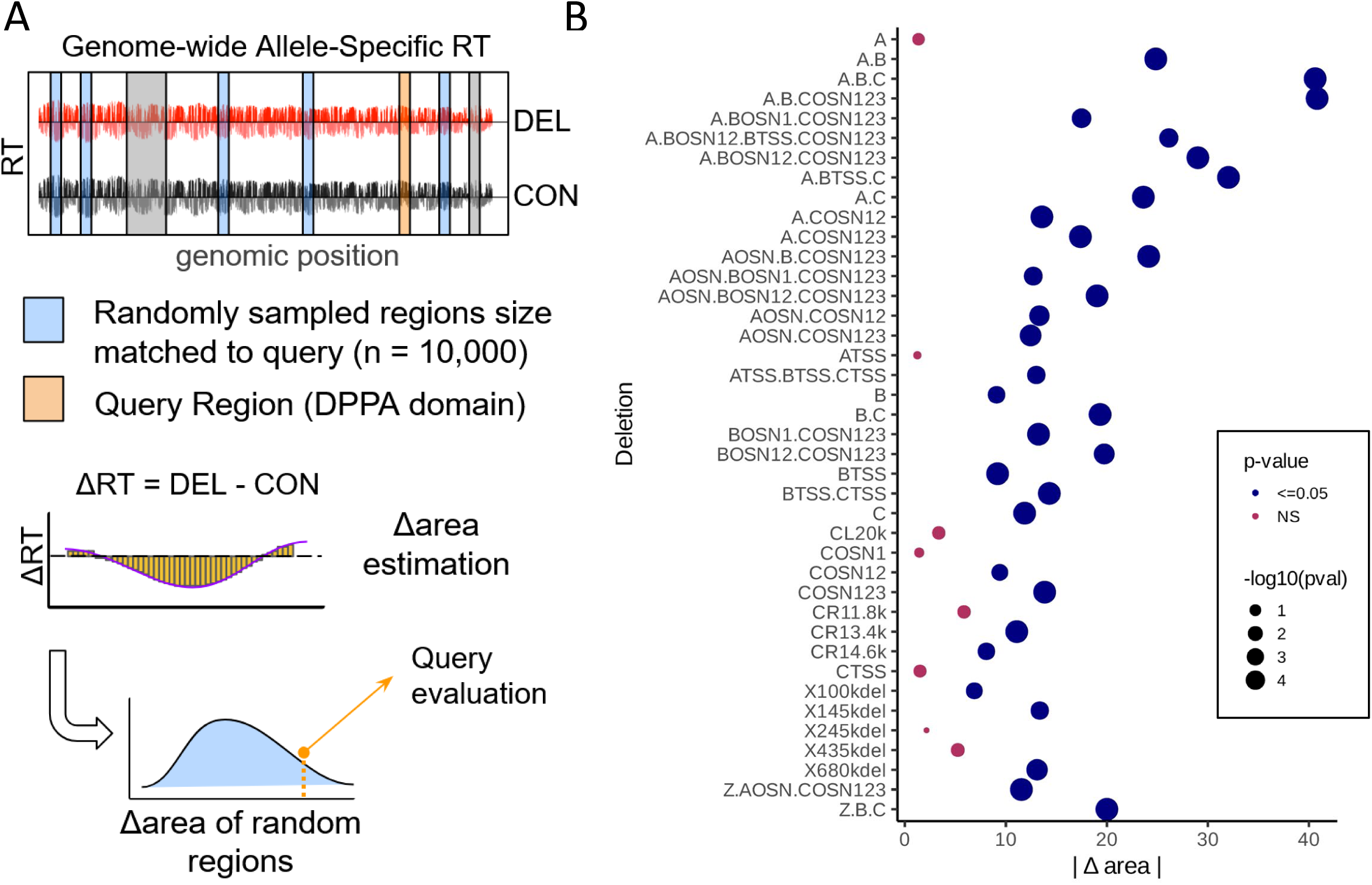
A) Schematic representation of AUC approach. B) |Δ area| for all deletions in this study with corresponding p-values. Color represents significance at p-value of 0.05 and size represents -log10(p-value).

**Supplemental Figure 3.**
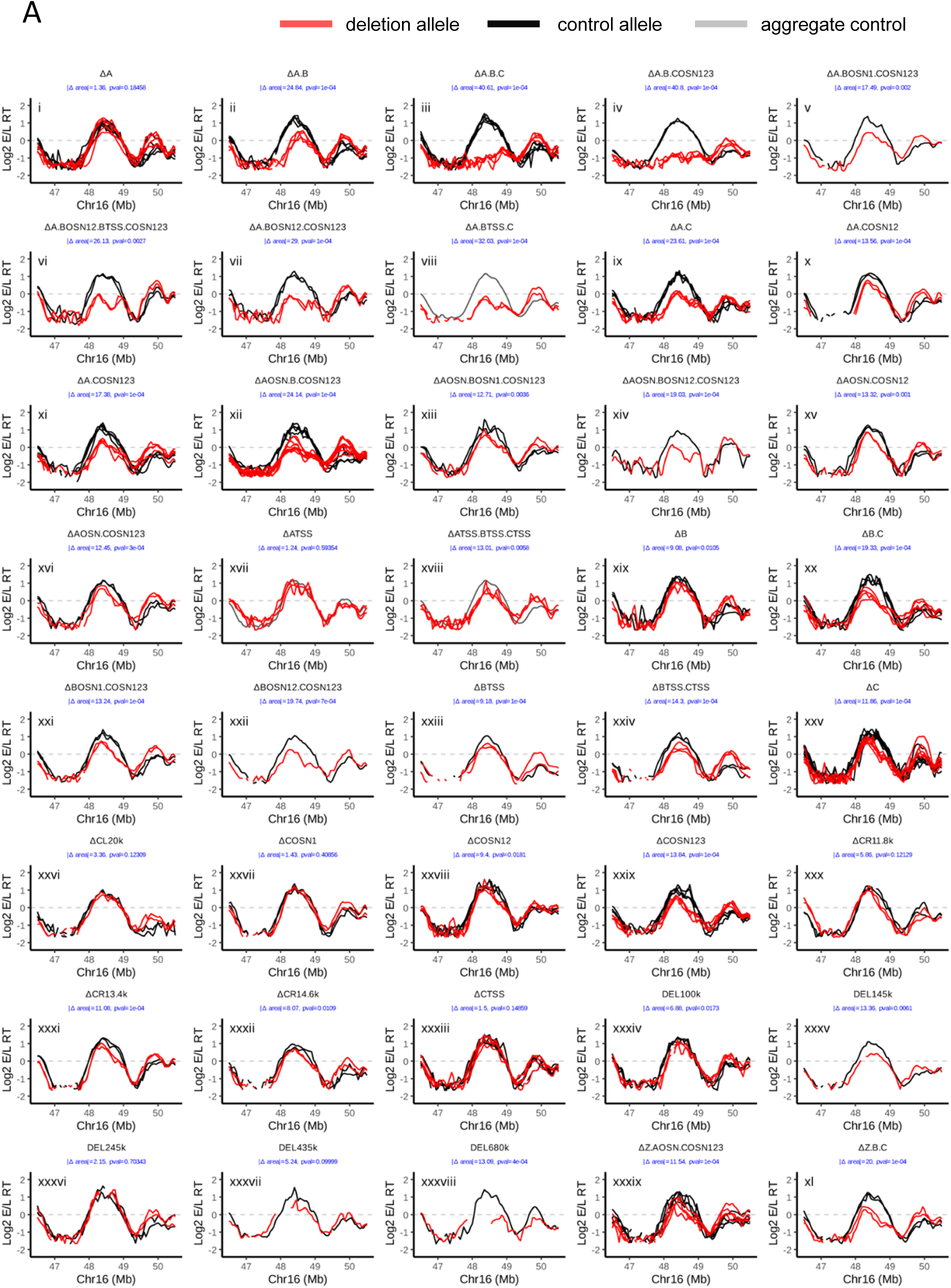
(i-xl) Allele specific E/L Repli-seq for all the data generated and analyzed in this study, including relevant deletions from our previous publication (Sima *et al*., 2019). Replicates (independent CRISPR-mediated deletion clones) are shown as different lines in each plot, comparing the deletion (red) allele to the homologous unmodified allele (black). In the occasional cases where a clone suffered one of the multiple deletions in both alleles we use an aggregated WT control (grey). The aggregated control was generated by averaging all RT profiles from the WT alleles across different deletions. The replication domain to the right of Dppa (c16: 49.65-50.05Mb mm10) shows some clonal variation in RT but there is a poor correlation (R = -0.35) between the effects of the deletions in the Dppa domain to this variation. Rather, this domain has been shown to display a genetic difference in RT between *m. castaneus* and *m. musculus* (Rivera-Mulia *et al*., 2018) as well as high cell to cell variability in RT (Zhao *et al*., 2020).

**Supplemental Figure 4.**
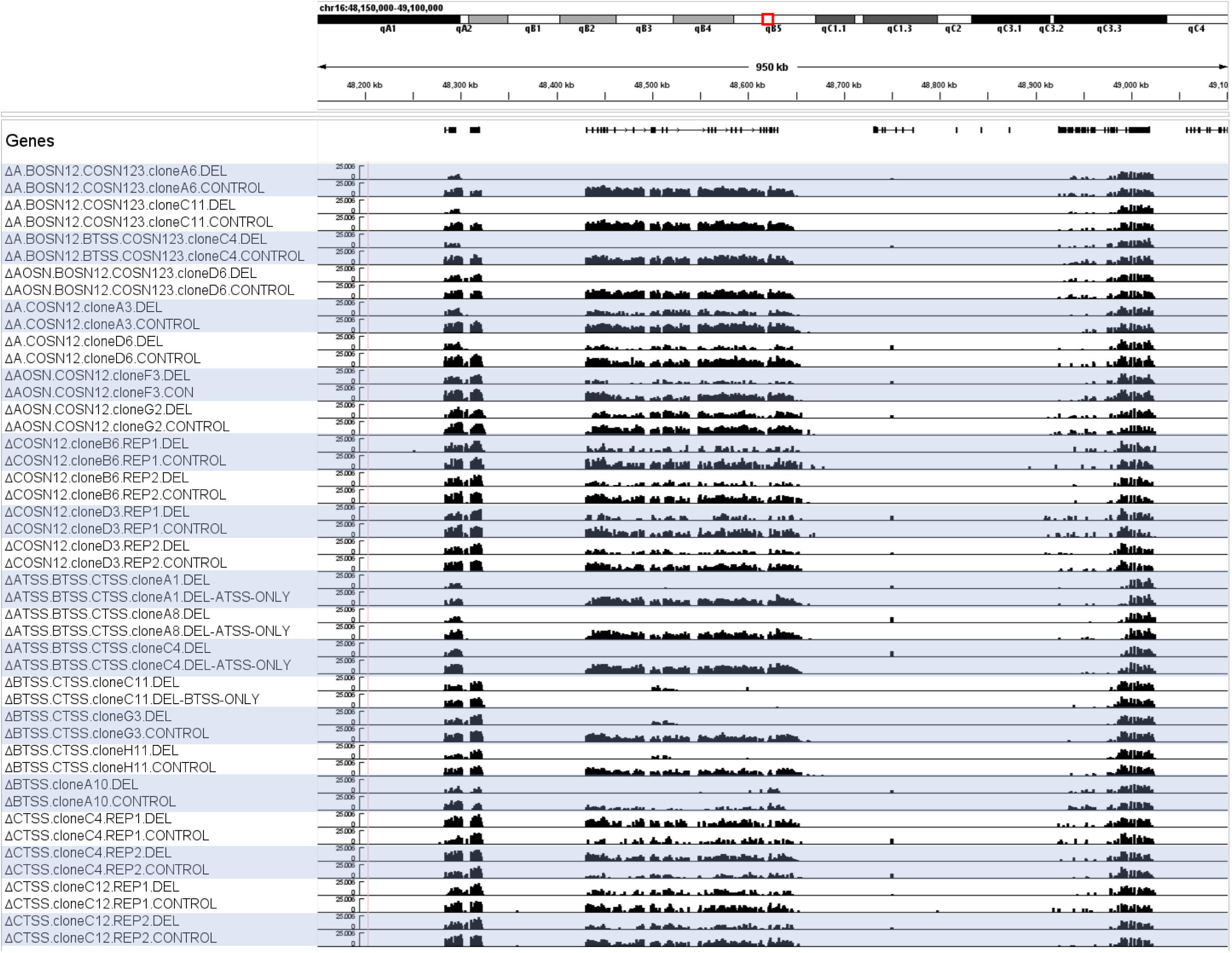
**All Bru-seq data sets generated in this study**. All of the BrU-seq generated in this study, displayed as in **Figure 4**. OSN deletion series data tracks are found on the top half while TSS deletion series data tracks can be found on the bottom half. We observed some variation in gene expression between the two alleles in WT cells (**Figure 4**). This variation can also be seen in the CTSS deletions (bottom tracks), as clone (C4) harbors a deletion of the *castaneus* allele whereas the other clone (C12) harbors a deletion on the *musculus* allele, and neither CTSS deletion has an effect on Morc1 expression.

**Fig S5.**
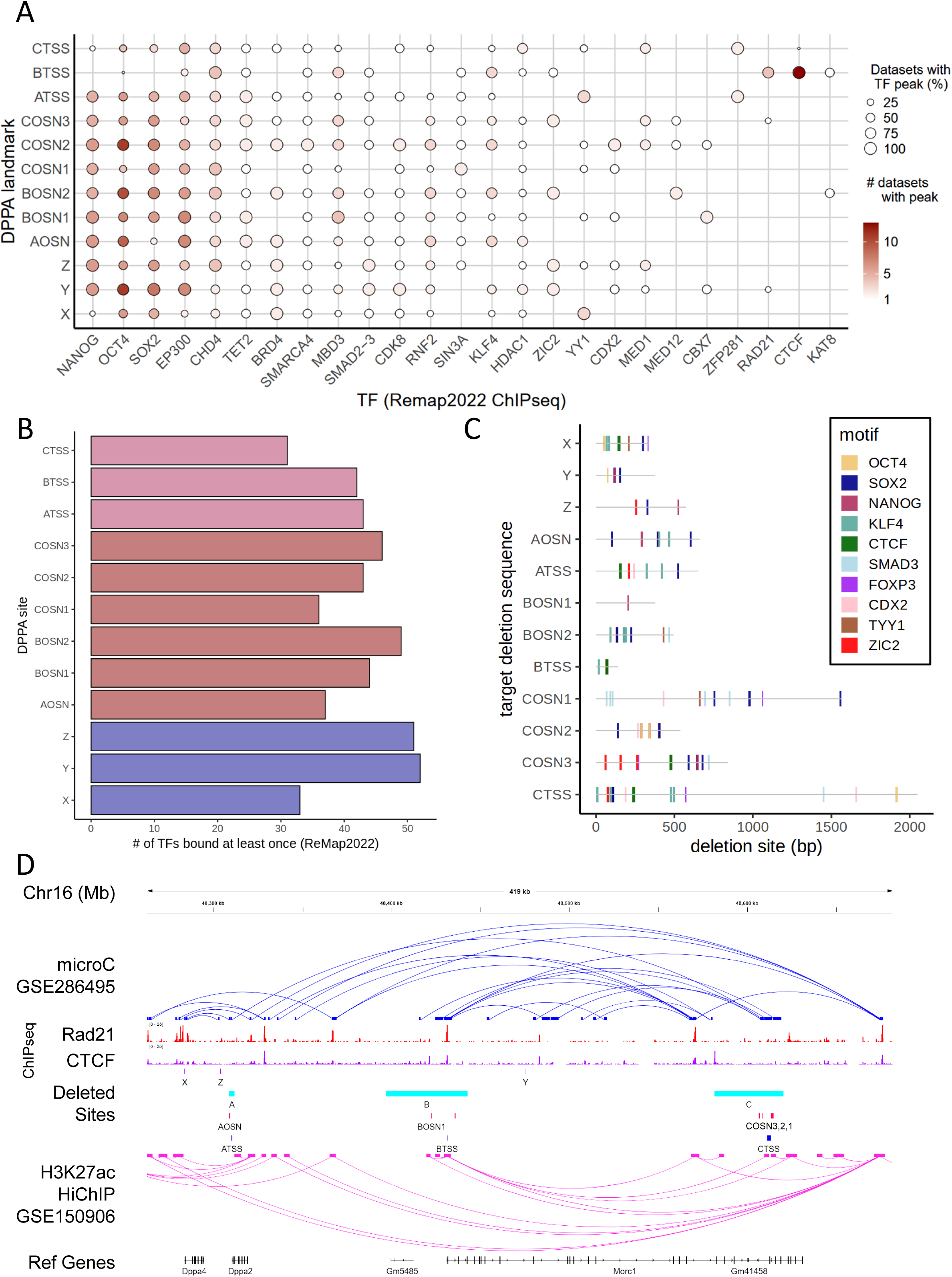
Deleted sites are diverse TFBS A) ChIPseq peaks from ‘mESC’ datasets in Remap 2022 overlapping at least two of the sites of interest in the DPPA domain. The size of the dots indicates the percentage of available ChIP datasets with a peak directly overlapping the site, and the color indicates the number of datasets with peak. B) TF binding diversity (Hammal *et al*., 2022) quantified as the number of TFs with at least one ChIPseq peak from Remap2022 ‘mESC’ overlapping the sites of interest. C) Putative TFBS in target deletion sequences determined by motif presence. Due to the presence of over 400 TF binding motifs in this collection of elements, only motifs for TFs from the ChIPseq targets shown on panel A are included. D) Micro-C (Jusuf *et al*., 2025) and H3K27ac HiChIP loops (Kraft *et al*., 2022) in the Dppa domain, as well as CTCF and Rad21 ChIPseq signal over input (Cattoglio *et al*., 2019; Hansen *et al*., 2017). Loop resolution varies between datasets and is indicated on y axis.

## SUPPLEMENTAL TABLE LEGENDS

**Supplemental Table 1 –** All sgRNA and primer sequences used to generate the deletions shown throughout this study.

**Supplemental Table 2 –** Chromosomal coordinates of all deletions shown in this study. This table also includes the sgRNA pairings used to generate all deletions.

**Supplemental Table 3 –** Genotypes of all cell lines generated in this study. Deletion genotypes (coordinates are available in **Supplemental Table 2**) were initially determined by PCR with the primers listed in **Supplemental Table 1**. Deletion PCR amplicons were then sanger sequenced to verify the breakpoints and the parental allele.

**Supplemental Table 4 –** Pairwise comparisons between relevant deletion pairs (ΔΔ area) and corresponding p-values.

**Supplemental Table 5 –** BruSeq Quantification.

## REFERENCES

1. Akerman I, Kasaai B, Bazarova A, Sang PB, Peiffer I, Artufel M, Derelle R, Smith G, Rodriguez-Martinez M, Romano M et al (2020) A predictable conserved DNA base composition signature defines human core DNA replication origins. Nat Commun 11: 4826

2. Bechhoefer J, Rhind N (2012) Replication timing and its emergence from stochastic processes. Trends Genet 28: 374–381

3. Besnard E, Babled A, Lapasset L, Milhavet O, Parrinello H, Dantec C, Marin JM, Lemaitre JM (2012) Unraveling cell type-specific and reprogrammable human replication origin signatures associated with G-quadruplex consensus motifs. Nat Struct Mol Biol 19: 837–844

4. Blayney JW, Francis H, Rampasekova A, Camellato B, Mitchell L, Stolper R, Cornell L, Babbs C, Boeke JD, Higgs DR et al (2023) Super-enhancers include classical enhancers and facilitators to fully activate gene expression. Cell 186: 5826–5839 e5818

5. Blin M, Le Tallec B, Nahse V, Schmidt M, Brossas C, Millot GA, Prioleau MN, Debatisse M (2019) Transcription-dependent regulation of replication dynamics modulates genome stability. Nat Struct Mol Biol 26: 58-66

6. Blumenfeld B, Ben-Zimra M, Simon I (2017) Perturbations in the Replication Program Contribute to Genomic Instability in Cancer. International Journal of Molecular Sciences 18: 1138

7. Bolstad B, 2023. preprocessCore: A collection of pre-processing functions. Bioconductor, p. Version 1.62.61.

8. Brossas C, Valton AL, Venev SV, Chilaka S, Counillon A, Laurent M, Goncalves C, Duriez B, Picard F, Dekker J et al (2020) Clustering of strong replicators associated with active promoters is sufficient to establish an early-replicating domain. EMBO J 39: e99520

9. Brueckner L, Zhao PA, Van Schaik T, Leemans C, Sima J, Peric-Hupkes D, Gilbert DM, Van Steensel B (2020) Local rewiring of genome–nuclear lamina interactions by transcription. The EMBO Journal 39

10. Carrington JT, Wilson RHC, Thiyagarajan S, Barker T, Catchpole L, Durrant A, Knitlhoffer V, Watkins C, Gharbi K, Nieduszynski CA (2024) Most human DNA replication initiation is dispersed throughout the genome with only a minority within previously identified initiation zones. bioRxiv: 2024.2004.2028.591325

11. Cattoglio C, Pustova I, Walther N, Ho JJ, Hantsche-Grininger M, Inouye CJ, Hossain MJ, Dailey GM, Ellenberg J, Darzacq X et al (2019) Determining cellular CTCF and cohesin abundances to constrain 3D genome models. Elife 8

12. Cayrou C, Ballester B, Peiffer I, Fenouil R, Coulombe P, Andrau J-C, Van Helden J, Méchali M (2015) The chromatin environment shapes DNA replication origin organization and defines origin classes. Genome Research 25: 1873–1885

13. Chen YH, Keegan S, Kahli M, Tonzi P, Fenyo D, Huang TT, Smith DJ (2019) Transcription shapes DNA replication initiation and termination in human cells. Nat Struct Mol Biol 26: 67–77

14. Choi K-J, Quan MD, Qi C, Lee J-H, Tsoi PS, Zahabiyon M, Bajic A, Hu L, Prasad BVV, Liao S-CJ et al (2022) NANOG prion-like assembly mediates DNA bridging to facilitate chromatin reorganization and activation of pluripotency. Nature Cell Biology 24: 737–747

15. de Wit E, Bouwman BA, Zhu Y, Klous P, Splinter E, Verstegen MJ, Krijger PH, Festuccia N, Nora EP, Welling M et al (2013) The pluripotent genome in three dimensions is shaped around pluripotency factors. Nature 501: 227–231

16. Di Giammartino DC, Kloetgen A, Polyzos A, Liu Y, Kim D, Murphy D, Abuhashem A, Cavaliere P, Aronson B, Shah V et al (2019) KLF4 is involved in the organization and regulation of pluripotency-associated three-dimensional enhancer networks. Nature Cell Biology 21: 1179–1190

17. Dileep V, Ay F, Sima J, Vera DL, Noble WS, Gilbert DM (2015) Topologically associating domains and their long-range contacts are established during early G1 coincident with the establishment of the replication-timing program. Genome Research 25: 1104–1113

18. Dimitrova DS, Gilbert DM (1999) The spatial position and replication timing of chromosomal domains are both established in early G1 phase. Mol Cell 4: 983–993

19. Dowen JM, Fan ZP, Hnisz D, Ren G, Abraham BJ, Zhang LN, Weintraub AS, Schujiers J, Lee TI, Zhao K et al (2014) Control of cell identity genes occurs in insulated neighborhoods in mammalian chromosomes. Cell 159: 374-387

20. Fang D, Lengronne A, Shi D, Forey R, Skrzypczak M, Ginalski K, Yan C, Wang X, Cao Q, Pasero P et al (2017) Dbf4 recruitment by forkhead transcription factors defines an upstream rate-limiting step in determining origin firing timing. Genes & Development 31: 2405–2415

21. Gabut M, Samavarchi-Tehrani P, Wang X, Slobodeniuc V, O’Hanlon D, Sung H-K, Alvarez M, Talukder S, Pan Q, Mazzoni O, Esteban et al (2011) An Alternative Splicing Switch Regulates Embryonic Stem Cell Pluripotency and Reprogramming. Cell 147: 132–146

22. Gerhardt J, Tomishima MJ, Zaninovic N, Colak D, Yan Z, Zhan Q, Rosenwaks Z, Jaffrey SR, Schildkraut CL (2014) The DNA replication program is altered at the FMR1 locus in fragile X embryonic stem cells. Mol Cell 53: 19–31

23. Gilbert DM (2001) Nuclear position leaves its mark on replication timing. J Cell Biol 152: F11–15

24. Goel VY, Aboreden NG, Jusuf JM, Zhang H, Mori LP, Mirny LA, Blobel GA, Banigan EJ, Hansen AS (2024) Dynamics of microcompartment formation at the mitosis-to-G1 transition. *bioRxiv*

25. Göös H, Kinnunen M, Salokas K, Tan Z, Liu X, Yadav L, Zhang Q, Wei G-H, Varjosalo M (2022) Human transcription factor protein interaction networks. Nature Communications 13

26. Grant CE, Bailey TL, Noble WS (2011) FIMO: scanning for occurrences of a given motif. Bioinformatics 27: 1017–1018

27. Guilbaud G, Murat P, Wilkes HS, Lerner LK, Sale JE, Krude T (2022) Determination of human DNA replication origin position and efficiency reveals principles of initiation zone organisation. Nucleic Acids Res 50: 7436–7450

28. Hammal F, de Langen P, Bergon A, Lopez F, Ballester B (2022) ReMap 2022: a database of Human, Mouse, Drosophila and Arabidopsis regulatory regions from an integrative analysis of DNA-binding sequencing experiments. Nucleic Acids Res 50: D316–D325

29. Hansen AS, Pustova I, Cattoglio C, Tjian R, Darzacq X (2017) CTCF and cohesin regulate chromatin loop stability with distinct dynamics. Elife 6

30. He Y, Hariharan M, Gorkin DU, Dickel DE, Luo C, Castanon RG, Nery JR, Lee AY, Zhao Y, Huang H et al (2020) Spatiotemporal DNA methylome dynamics of the developing mouse fetus. Nature 583: 752–759

31. Heskett MB, Smith LG, Spellman P, Thayer MJ (2020) Reciprocal monoallelic expression of ASAR lncRNA genes controls replication timing of human chromosome 6. RNA 26: 724–738

32. Hiratani I, Ryba T, Itoh M, Rathjen J, Kulik M, Papp B, Fussner E, Bazett-Jones DP, Plath K, Dalton S et al (2010) Genome-wide dynamics of replication timing revealed by in vitro models of mouse embryogenesis. Genome Research 20: 155–169

33. Hiratani I, Ryba T, Itoh M, Yokochi T, Schwaiger M, Chang C-W, Lyou Y, Townes TM, Schübeler D, Gilbert DM (2008) Global Reorganization of Replication Domains During Embryonic Stem Cell Differentiation. PLoS Biology 6: e245

34. Hu Y, Salgado Figueroa D, Zhang Z, Veselits M, Bhattacharyya S, Kashiwagi M, Clark MR, Morgan BA, Ay F, Georgopoulos K (2023) Lineage-specific 3D genome organization is assembled at multiple scales by IKAROS. Cell 186: 5269–5289 e5222

35. Hu Y, Stillman B (2023) Origins of DNA replication in eukaryotes. Mol Cell 83: 352-372

36. Hyrien O (2015) Peaks cloaked in the mist: the landscape of mammalian replication origins. J Cell Biol 208: 147-160

37. Inoue N, Hess KD, Moreadith RW, Richardson LL, Handel MA, Watson ML, Zinn AR (1999) New gene family defined by MORC, a nuclear protein required for mouse spermatogenesis. Hum Mol Genet 8: 1201–1207

38. Jodkowska K, Pancaldi V, Rigau M, Almeida R, Fernandez-Justel JM, Grana-Castro O, Rodriguez-Acebes S, Rubio-Camarillo M, Carrillo-de Santa Pau E, Pisano D et al (2022) 3D chromatin connectivity underlies replication origin efficiency in mouse embryonic stem cells. Nucleic Acids Res 50: 12149–12165

39. Jusuf JM, Grosse-Holz S, Gabriele M, Mach P, Flyamer IM, Zechner C, Giorgetti L, Mirny L, Hansen AS (2025) Genome-wide absolute quantification of chromatin looping. bioRxiv

40. Khan A, Zhang X (2016) dbSUPER: a database of super-enhancers in mouse and human genome. Nucleic Acids Res 44: D164–171

41. King HW, Klose RJ (2017) The pioneer factor OCT4 requires the chromatin remodeller BRG1 to support gene regulatory element function in mouse embryonic stem cells. eLife 6

42. Klein KN, Zhao PA, Lyu X, Sasaki T, Bartlett DA, Singh AM, Tasan I, Zhang M, Watts LP, Hiraga SI et al (2021) Replication timing maintains the global epigenetic state in human cells. Science 372: 371–378

43. Knott RV, Simon, Peace M, Jared, Ostrow Z, A., Gan Y, Rex E, Alexandra, Viggiani J, Christopher, Tavaré S, Aparicio M, Oscar (2012) Forkhead Transcription Factors Establish Origin Timing and Long-Range Clustering in S. cerevisiae. Cell 148: 99–111

44. Kraft K, Yost KE, Murphy SE, Magg A, Long Y, Corces MR, Granja JM, Wittler L, Mundlos S, Cech TR et al (2022) Polycomb-mediated genome architecture enables long-range spreading of H3K27 methylation. Proc Natl Acad Sci U S A 119: e2201883119

45. Krueger F, Andrews SR (2016) SNPsplit: Allele-specific splitting of alignments between genomes with known SNP genotypes. F1000Res 5: 1479

46. Kulakovskiy IV, Vorontsov IE, Yevshin IS, Sharipov RN, Fedorova AD, Rumynskiy EI, Medvedeva YA, Magana-Mora A, Bajic VB, Papatsenko DA et al (2018) HOCOMOCO: towards a complete collection of transcription factor binding models for human and mouse via large-scale ChIP-Seq analysis. Nucleic Acids Res 46: D252–D259

47. Langmead B, Salzberg SL (2012) Fast gapped-read alignment with Bowtie 2. Nature Methods 9: 357–359

48. Li H, Handsaker B, Wysoker A, Fennell T, Ruan J, Homer N, Marth G, Abecasis G, Durbin R, Subgroup GPDP (2009) The Sequence Alignment/Map format and SAMtools. Bioinformatics 25: 2078–2079

49. Lichauco C, Foss EJ, Gatbonton-Schwager T, Athow NF, Lofts B, Acob R, Taylor E, Marquez JJ, Lao U, Miles S et al (2025) Sir2 and Fun30 regulate ribosomal DNA replication timing via MCM helicase positioning and nucleosome occupancy. Elife 13

50. Liu Y, Ai C, Gan T, Wu J, Jiang Y, Liu X, Lu R, Gao N, Li Q, Ji X et al (2021) Transcription shapes DNA replication initiation to preserve genome integrity. Genome Biol 22: 176

51. Liu Y, Wan X, Li H, Chen Y, Hu X, Chen H, Zhu D, Li C, Zhang Y (2023) CTCF coordinates cell fate specification via orchestrating regulatory hubs with pioneer transcription factors. Cell Rep 42: 113259

52. Lo JH-H, Edwards M, Langerman J, Sridharan R, Plath K, Smale ST (2022) Oct4:Sox2 binding is essential for establishing but not maintaining active and silent states of dynamically regulated genes in pluripotent cells. Genes & Development 36: 1079–1095

53. Madsen JGS, Madsen MS, Rauch A, Traynor S, Van Hauwaert EL, Haakonsson AK, Javierre BM, Hyldahl M, Fraser P, Mandrup S (2020) Highly interconnected enhancer communities control lineage-determining genes in human mesenchymal stem cells. Nat Genet 52: 1227-1238

54. Malzl D, Peycheva M, Rahjouei A, Gnan S, Klein KN, Nazarova M, Schoeberl UE, Gilbert DM, Buonomo SCB, Virgilio MD et al, 2023. RIF1 regulates replication origin activity and early replication timing in B cells. Cold Spring Harbor Laboratory.

55. Marchal C, Sasaki T, Vera D, Wilson K, Sima J, Rivera-Mulia JC, Trevilla-García C, Nogues C, Nafie E, Gilbert DM (2018) Genome-wide analysis of replication timing by next-generation sequencing with E/L Repli-seq. Nat Protoc 13: 819–839

56. Martin M (2011) Cutadapt removes adapter sequences from high-throughput sequencing reads. EMBnetjournal 17: 10

57. Noguchi S, Arakawa T, Fukuda S, Furuno M, Hasegawa A, Hori F, Ishikawa-Kato S, Kaida K, Kaiho A, Kanamori-Katayama M et al (2017) FANTOM5 CAGE profiles of human and mouse samples. Sci Data 4: 170112

58. Ostrow AZ, Kalhor R, Gan Y, Villwock SK, Linke C, Barberis M, Chen L, Aparicio OM (2017) Conserved forkhead dimerization motif controls DNA replication timing and spatial organization of chromosomes in *S. cerevisiae*. Proceedings of the National Academy of Sciences 114: E2411–E2419

59. Paulsen MT, Veloso A, Prasad J, Bedi K, Ljungman EA, Magnuson B, Wilson TE, Ljungman M (2014) Use of Bru-Seq and BruChase-Seq for genome-wide assessment of the synthesis and stability of RNA. Methods 67: 45–54

60. Petryk N, Dalby M, Wenger A, Stromme CB, Strandsby A, Andersson R, Groth A (2018) MCM2 promotes symmetric inheritance of modified histones during DNA replication. Science 361: 1389–1392

61. Petryk N, Kahli M, d’Aubenton-Carafa Y, Jaszczyszyn Y, Shen Y, Silvain M, Thermes C, Chen CL, Hyrien O (2016) Replication landscape of the human genome. Nat Commun 7: 10208

62. Platt EJ, Smith L, Thayer MJ (2018) L1 retrotransposon antisense RNA within ASAR lncRNAs controls chromosome-wide replication timing. Journal of Cell Biology 217: 541–553

63. Pratto F, Brick K, Cheng G, Lam KG, Cloutier JM, Dahiya D, Wellard SR, Jordan PW, Camerini-Otero RD (2021) Meiotic recombination mirrors patterns of germline replication in mice and humans. Cell 184: 4251–4267 e4220

64. Prorok P, Artufel M, Aze A, Coulombe P, Peiffer I, Lacroix L, Guédin A, Mergny J-L, Damaschke J, Schepers A et al (2019) Involvement of G-quadruplex regions in mammalian replication origin activity. Nature Communications 10

65. Quinlan AR, Hall IM (2010) BEDTools: a flexible suite of utilities for comparing genomic features. Bioinformatics 26: 841–842

66. Reyna J, Fetter K, Ignacio R, Marandi CCA, Rao N, Jiang Z, Figueroa DS, Bhattacharyya S, Ay F (2024) Loop Catalog: a comprehensive HiChIP database of human and mouse samples. *bioRxiv*

67. Rivera-Mulia C, Juan, Gilbert M, David (2016) Replicating Large Genomes: Divide and Conquer. Molecular Cell 62: 756–765

68. Rivera-Mulia JC, Desprat R, Trevilla-Garcia C, Cornacchia D, Schwerer H, Sasaki T, Sima J, Fells T, Studer L, Lemaitre J-M et al (2017) DNA replication timing alterations identify common markers between distinct progeroid diseases. Proceedings of the National Academy of Sciences 114: E10972–E10980

69. Rivera-Mulia JC, Dimond A, Vera D, Trevilla-Garcia C, Sasaki T, Zimmerman J, Dupont C, Gribnau J, Fraser P, Gilbert DM (2018) Allele-specific control of replication timing and genome organization during development. Genome Research 28: 800–811

70. Rivera-Mulia JC, Kim S, Gabr H, Chakraborty A, Ay F, Kahveci T, Gilbert DM (2019a) Replication timing networks reveal a link between transcription regulatory circuits and replication timing control. Genome Research 29: 1415–1428

71. Rivera-Mulia JC, Sasaki T, Trevilla-Garcia C, Nakamichi N, Knapp D, Hammond CA, Chang BH, Tyner JW, Devidas M, Zimmerman J et al (2019b) Replication timing alterations in leukemia affect clinically relevant chromosome domains. Blood Adv 3: 3201–3213

72. Rothenberg EV (2022) Transcription factors specifically control change. Genes & Development 36: 1097–1099

73. Sasaki T, Ramanathan S, Okuno Y, Kumagai C, Shaikh SS, Gilbert DM (2006) The Chinese hamster dihydrofolate reductase replication origin decision point follows activation of transcription and suppresses initiation of replication within transcription units. Mol Cell Biol 26: 1051–1062

74. Sasaki T, Rivera-Mulia JC, Vera D, Zimmerman J, Das S, Padget M, Nakamichi N, Chang BH, Tyner J, Druker BJ et al (2017) Stability of patient-specific features of altered DNA replication timing in xenografts of primary human acute lymphoblastic leukemia. Exp Hematol 51: 71–82.e73

75. Scherr MJ, Wahab SA, Remus D, Duderstadt KE (2022) Mobile origin-licensing factors confer resistance to conflicts with RNA polymerase. Cell Rep 38: 110531

76. Sima J, Chakraborty A, Dileep V, Michalski M, Klein KN, Holcomb NP, Turner JL, Paulsen MT, Rivera-Mulia JC, Trevilla-Garcia C et al (2019) Identifying cis Elements for Spatiotemporal Control of Mammalian DNA Replication. Cell 176: 816–830.e818

77. Singh G, Mullany S, Moorthy SD, Zhang R, Mehdi T, Tian R, Duncan AG, Moses AM, Mitchell JA (2021) A flexible repertoire of transcription factor binding sites and a diversity threshold determines enhancer activity in embryonic stem cells. Genome Res 31: 564–575

78. Takebayashi S-I, Dileep V, Ryba T, Dennis JH, Gilbert DM (2012a) Chromatin-interaction compartment switch at developmentally regulated chromosomal domains reveals an unusual principle of chromatin folding. Proceedings of the National Academy of Sciences 109: 12574–12579

79. Takebayashi S-I, Ryba T, Gilbert DM (2012b) Developmental control of replication timing defines a new breed of chromosomal domains with a novel mechanism of chromatin unfolding. Nucleus 3: 500–507

80. Therizols P, Illingworth RS, Courilleau C, Boyle S, Wood AJ, Bickmore WA (2014) Chromatin decondensation is sufficient to alter nuclear organization in embryonic stem cells. Science 346: 1238–1242

81. Vouzas A, Sasaki T, Rivera-Mulia C, Juan, Turner JL, Brown AN, Alexander KE, Brueckner L, van Steensel B, Gilbert DM (2025) Transcription can be sufficient, but is not necessary, to advance replication timing. *bioRxiv*

82. Vouzas AE, Gilbert DM (2023) Replication timing and transcriptional control: beyond cause and effect - part IV. Curr Opin Genet Dev 79: 102031

83. Wang R, Chen F, Chen Q, Wan X, Shi M, Chen AK, Ma Z, Li G, Wang M, Ying Y et al (2022a) MyoD is a 3D genome structure organizer for muscle cell identity. Nat Commun 13: 205

84. Wang W, Chandra A, Goldman N, Yoon S, Ferrari EK, Nguyen SC, Joyce EF, Vahedi G (2022b) TCF-1 promotes chromatin interactions across topologically associating domains in T cell progenitors. Nat Immunol 23: 1052–1062

85. Wang W, Klein KN, Proesmans K, Yang H, Marchal C, Zhu X, Borrman T, Hastie A, Weng Z, Bechhoefer J et al (2021) Genome-wide mapping of human DNA replication by optical replication mapping supports a stochastic model of eukaryotic replication. Mol Cell 81: 2975–2988 e2976

86. Winick-Ng W, Kukalev A, Harabula I, Zea-Redondo L, Szabo D, Meijer M, Serebreni L, Zhang Y, Bianco S, Chiariello AM et al (2021) Cell-type specialization is encoded by specific chromatin topologies. Nature 599: 684–691

87. Zhao PA, Sasaki T, Gilbert DM (2020) High-resolution Repli-Seq defines the temporal choreography of initiation, elongation and termination of replication in mammalian cells. Genome Biology 21

